# The cingulo-opercular network is composed of two distinct sub-systems

**DOI:** 10.1101/2022.09.16.508254

**Authors:** Caterina Gratton, Ally Dworetsky, Babatunde Adeyemo, Benjamin A. Seitzman, Derek M. Smith, Steven E. Petersen, Maital Neta

## Abstract

The cingulo-opercular (CO) network and its two best studied regions – the dorsal anterior cingulate and anterior insula – have been linked to task control, but also implicated in many additional processes across cognitive, social, and emotional domains. However, most prior work investigating the CO network has used a group-average approach, which may mix signals across nearby regions that vary across individuals. Here, we reevaluate the CO network’s role in task control with both task and rest fMRI, using regions with a high probability of CO network agreement across individuals. Hierarchical clustering analyses suggest heterogeneity in the CO network’s task response properties, with one sub-system (CO1) showing consistency with prior task control characterizations while another sub-system (CO2) has weak task control responses, but preserved ties to pain and motor functions. Resting-state connectivity confirms subtle differences in the architecture of these two sub-systems. This evidence suggests that, when individual variation in network locations is addressed, the CO network includes (at least) two linked sub-systems with differential roles in task control and other cognitive/motor/interoceptive responses, which may help explain varied accounts of its functions. We propose that this fractionation may reflect expansion of primary CO body-oriented control functions to broader domain-general contexts.

## INTRODUCTION

There is an ever-growing body of literature that cuts across many domains of inquiry that has explored the functional role of the cingulo-opercular (CO) brain network. CO network regions include the dorsal anterior cingulate and medial superior frontal cortex (dACC/msFC)^1^, along with the anterior insula, sometimes spreading to the frontal operculum (aI/fO). This network shows strong spontaneous correlations at rest (Dosenbach et al. 2007; Power et al. 2011) and is among the most commonly activated across a wide range of neuroimaging tasks (**Fig. 1A**; (Nelson et al. 2010)).

**Figure 1:**
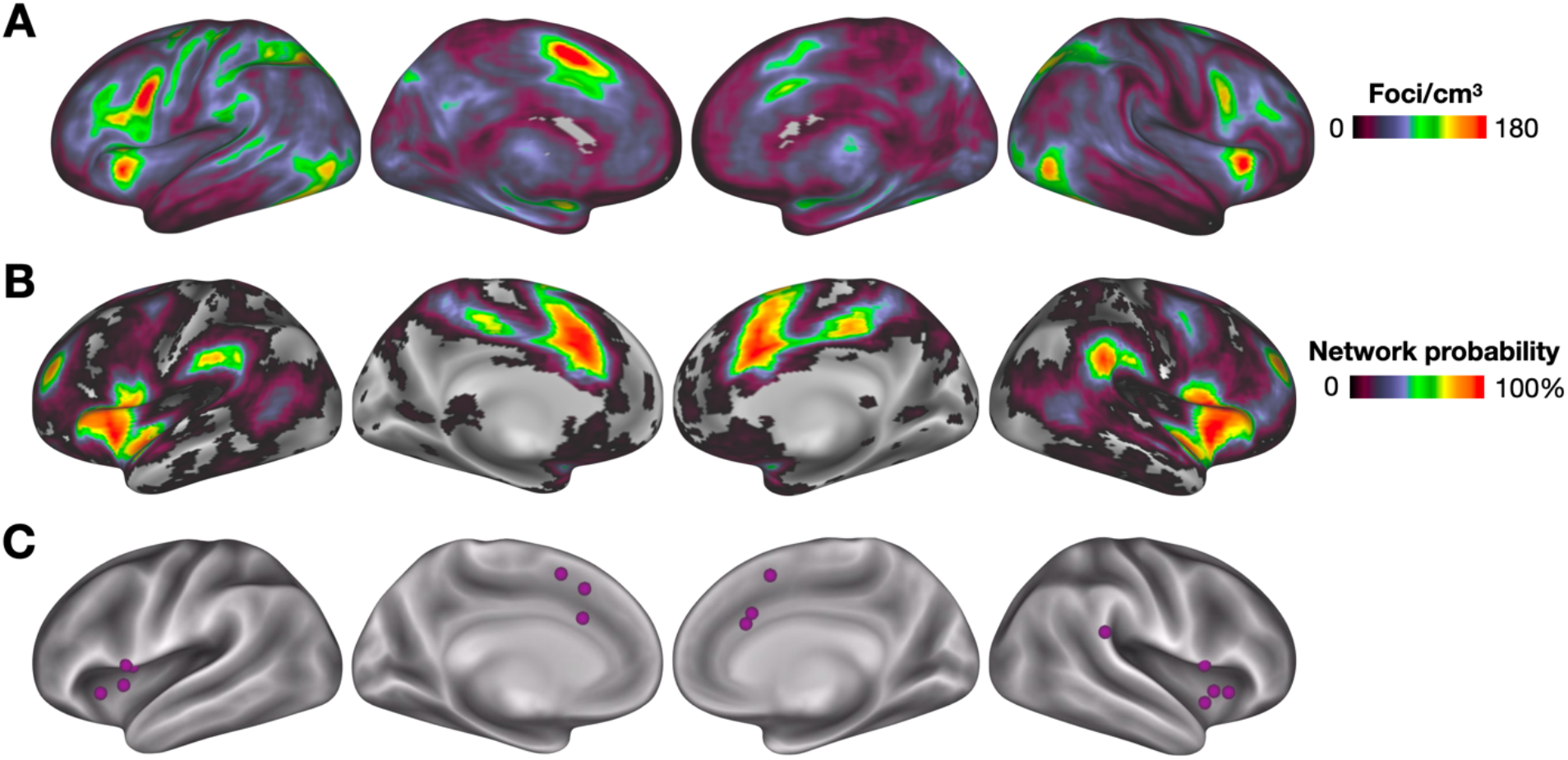
**(A)** A consensus map of task-related activation across over 1,000 brain imaging studies; warmer colors indicate locations that are more consistently activated across studies. **(B)** A probabilistic map of the cingulo-opercular network defined via resting-state correlations; higher values indicate locations consistently identified as cingulo-opercular across individuals (Dworetsky et al. 2021). **(C)** From this probabilistic map, 15 regions of interest were selected from Power et al. (2011) that have a high probability (>75%) of being identified as part of the CO network across participants.

Within our own work, we have suggested that the CO network has a broad role in task control, or the ability of humans to easily and flexibly configure the specific processes necessary to perform many different tasks based on their goals (Dosenbach et al. 2008; Dosenbach et al. 2006; Gratton et al. 2018b; Power and Petersen 2013). This characterization is based on observations that regions in the CO network (especially dACC and aI/fO) are associated with a variety of control-related signals that play a role in initiating, maintaining, and adjusting to task demands, including start and stop cues (Dosenbach et al. 2006), sustained task signals (Dosenbach et al. 2006; Petersen and Dubis 2012), and signals relating to errors (Dosenbach et al. 2006; Neta et al. 2015), ambiguity (Neta et al. 2017; Neta et al. 2014), and other performance monitoring signals (Gratton et al. 2017); **Fig. 2**). However, additional studies to date have offered a range of explanations for the functional role of these regions, particularly the dACC, within many domains of cognitive, social, and affective neuroscience (Alexander and Brown 2011; Bartels and Zeki 2004; Brown and Braver 2007; Craig 2009; Grinband et al. 2011; Lieberman and Eisenberger 2015; Neta et al. 2013; Ochsner et al. 2009; Sadaghiani and D’Esposito 2015; Sterzer et al. 2002; Stolier and Freeman 2017; Thompson-Schill et al. 1997; Wager et al. 2009; Wessel et al. 2012).

**Figure 2:**
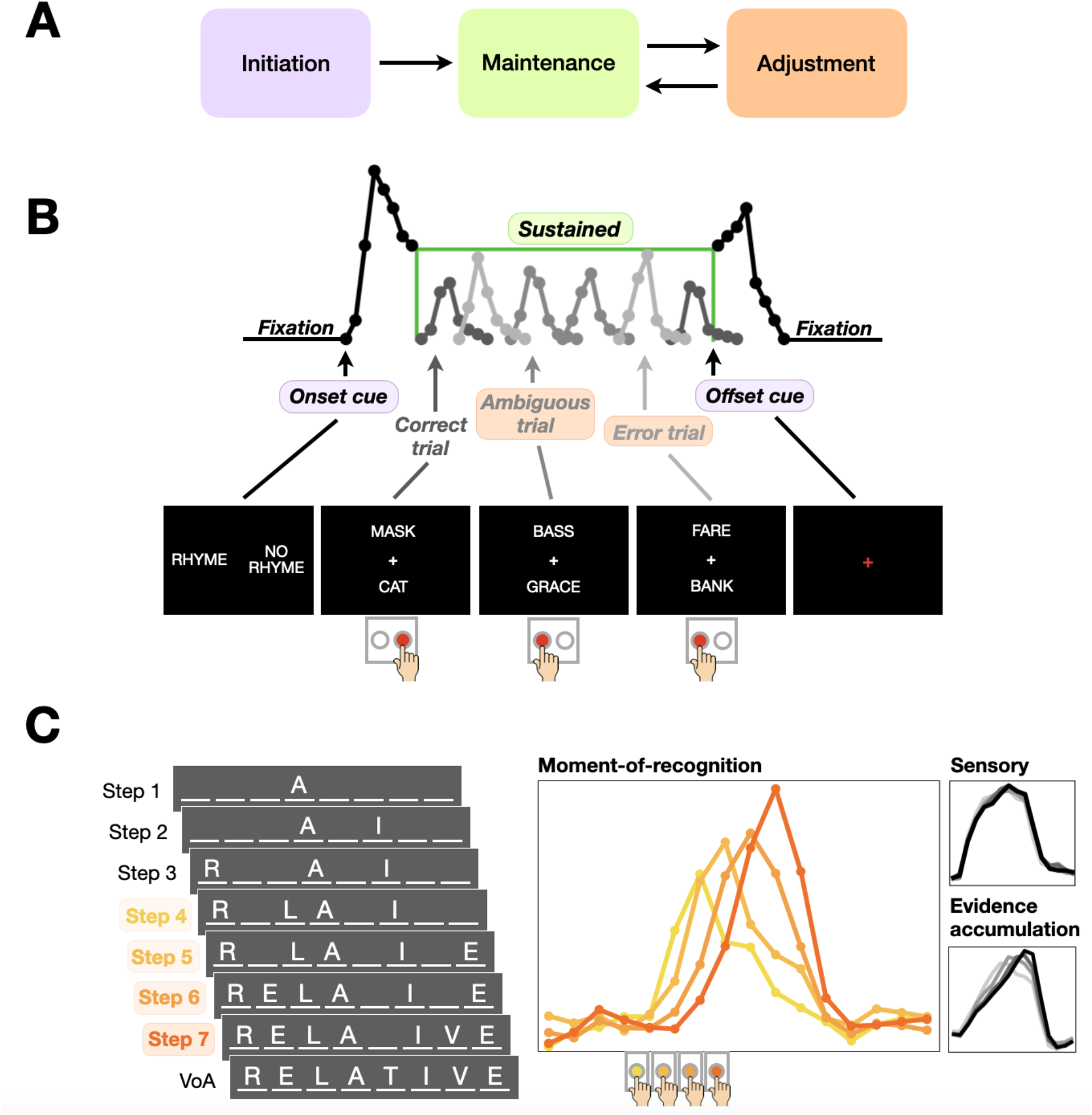
Control signal schematic. **(A)** In this work, we focused on analyzing task control signals associated with task set initiation, maintenance, and adjustment. These signals were extracted from mixed block/event-related designs (**B**; Tasks 1-16 in **Supp. Table 1**) and decision-making ‘slow reveal’ paradigms (**C**; Tasks 17-21 **in Supp. Table 1**). **(B)** Signals extracted from mixed block/event-related studies include: onset/offset cues at the start and end of each task block; signals from trials with an ambiguous response and trials with an error relative to correct trials; and a sustained activation over the entirety of the task block. In this example from a rhyme/no-rhyme task (e.g., Task 2), the onset cue is signaled by a display showing “rhyme” on the left side and “no rhyme” on the right side, indicating the button participants should press when shown two words that rhyme or do not rhyme. Sustained signals were extracted over the task block, and the offset cue was indicated by a red fixation cross. **(C)** The slow reveal tasks included examples of stimuli gradually revealed over the course of a trial (e.g., letters from a word, objects from behind a noise mask). Participants’ goal is to identify the final target item as quickly as possible, and verify that response at the end (VoA). Timecourses are extracted from these slow reveal tasks dependent on when a participant indicates recognition (e.g., at response step 4, 5, etc.; shaded colors). An illustration of the transient response timecourses over the course of the trial is shown in the middle for a canonical performance-related region; separate timecourses for trials with a button press from response steps 4 (earlier) to 7 (later) are represented from light to dark orange. Example timecourses at the far right insets depict other timecourse patterns associated with sensory processing (similar timecourse regardless of when the response is made) and evidence accumulation (early onsets with gradual, delayed peaks associated with the final response time).

One potential explanation for the broad range of functions ascribed to the CO network is heterogeneity in functional network organization (Smith et al. 2021). Indeed, a number of recent findings have highlighted that the spatial layout of functional networks varies across individuals, especially in frontal and parietal regions of the brain (Finn et al. 2015; Gratton et al. 2018a; Langs et al. 2016; Mueller et al. 2013; Seitzman et al. 2019). In contrast, most studies that have used tasks to probe the functions of the CO network – including our own – use standard group approaches (Dubis et al. 2016; Gratton et al. 2016; Neta et al. 2017; Neta et al. 2014; Sadaghiani and D’Esposito 2015; Sestieri et al. 2014; Vaden et al. 2013). These approaches average data together assuming a point-by-point spatial correspondence across individuals.

Individual variability in network organization means that these studies are likely mixing together signals from diverse networks across individuals, muddying interpretation of the underlying function(s) of these regions.

Within each of the large swaths of cortex associated with the CO system (e.g., dACC/msFC, aI/fO), it appears that there are multiple separate but closely adjacent networks (Power et al. 2011; Smith et al. 2021; Yeo et al. 2011). Precise definitions are needed to segregate the CO from its nearby networks (e.g., see dACC analysis in (Smith et al. 2021)). In addition, whole-brain functional connectivity analyses (e.g., (Gordon et al. 2017; Power et al. 2011) suggest that the CO network extends beyond the dACC and aI/fO to include posterior insular and supramarginal regions, as well as regions along the medial superior frontal and parietal cortex, and a region in the anterior dorsolateral prefrontal cortex (**Fig. 1B**). However, many analyses of the CO network restrict themselves to one or a small subset of these regions, making it unclear whether task responses are specific to sub-regions of the CO or if they extend throughout the full network.

While a substantial amount of resting-state data is necessary to achieve reliable whole-brain network definitions on an individual level (Elliott et al. 2019; Gordon et al. 2017; Laumann et al. 2015; Noble et al. 2017), we have recently developed a probabilistic atlas of functional networks that can be used to focus group analyses on locations with high consensus in network definitions across individuals (Dworetsky et al. 2021). This probabilistic atlas was used to define 153 whole-brain regions, including 15 regions of the CO network, with >75% consensus across individuals (**Fig. 1C**), a pattern validated across 4 diverse neuroimaging datasets. These high consensus CO regions can then be used to probe the properties of the CO network in group analyses, with high spatial precision, while reducing issues of heterogeneity in network definition across participants.

The goal of the present work was to explore our original characterization of the CO network – both through task and rest fMRI data – with higher precision in our network definitions that addresses variability in functional network locations across individuals. Specifically, we will use these high consensus CO regions to take a detailed look at the CO network and quantify the degree to which different regions of this system show common or distinct responses associated with task control. In addition, we will leverage resting-state patterns to examine the network architecture among these regions. This approach affords us the opportunity to re-examine the unity of CO region functions and place this view within the broader context of large-scale cortical systems. Given the ubiquitous nature of activity in these regions across many different domains of inquiry, a detailed decomposition of their functional activations (during tasks) and connectivity (at rest) would be a useful step toward advancing these disparate fields.

## METHODS

### Overview and datasets

The goal of this work was to re-evaluate previous results regarding the properties of the CO network, with a focus on regions showing a high consensus across individuals. To this end, we conducted a series of task and resting-state fMRI analyses with these regions. The task fMRI analyses include data from a total of 16 mixed block/event-related experiments and 5 experiments with a slow reveal (event-related) design, spanning a total of 421 human participants, previously collected in the Petersen-Schlaggar labs at Washington University in St. Louis (see **Supp. Table 1** for details on all experiments). Tasks were selected for inclusion in the meta-analysis based on their links to task control (Dosenbach et al. 2007; Dosenbach et al. 2006; Dubis et al. 2016) and decision-making (Ploran et al. 2007; Wheeler et al. 2008), processes that are hypothesized as central to the CO network.

The CO network was further interrogated through functional connectivity analyses using a resting-state fMRI dataset collected at Dartmouth College (N=69; (Gordon et al. 2016)). Subjects were healthy young adults between 18 and 35 years of age and ranged from 20 to 49 minutes of low motion rest data.

All participants provided informed consent and procedures were approved by the Washington University (for task meta-analyses) and Dartmouth (for resting state) Institutional Review Boards.

### fMRI tasks

Task details are outlined in **Supp. Fig 1**. A more detailed description of each task can be found in its respective original publication referenced in **Supp. Table 1**. In the present analyses, 16 mixed block/event-related and 5 slow reveal (event-related) tasks were included. In mixed block/event-related designs (Petersen and Dubis 2012), a cue was presented to signal the start of a task, followed by a task block period during which jittered trials were presented for each task (see schematic in **Fig. 2B**). At the end of the block, a cue was presented again to indicate the end of the task. Tasks varied in cognitive process (e.g., semantic, phonological, visual attention) and input/output modality and form, with stimuli presented across tasks including words, tones, images, dot pairs, and Gabor patches (**Supp. Table 1**). Mixed design tasks were used to measure task maintenance (sustained block signals from Tasks 1-16 in **Supp. Table 1**) as well as event-related responses to task initiation and adjustment (cues, errors, and ambiguous stimuli from Tasks 1-2 in **Supp. Table 1)**.

In the slow reveal event-related designs (Gratton et al. 2017; Ploran et al. 2007; Wheeler et al. 2008), a target item was hidden at the start of the trial and slowly revealed over the course of the trial (see example in **Fig. 2C**). Depending on the task, targets included object images or words, and participants were asked to either identify the targets or to determine whether they had previously been seen (Tasks 17-21, **Supp. Table 1**). Slow reveal tasks were used to estimate trial-level activity involved in decision-making.

### Image acquisition

Details on MRI acquisition parameters for all tasks included in the meta-analysis can be found in **Supp. Table 2**; briefly, data from tasks run at varying scanner strengths were included (1.5T or 3T), at TRs ranging from 2s to 3.18s, and at varying voxel resolutions (4mm isotropic, 3.2mm isotropic, 3.75×3.75×8mm, or 3×3×3.5mm). For all functional runs, whole-brain EPI acquisitions (MR frames) were obtained using a spin-echo or gradient-echo BOLD contrast sensitive sequence and acquired parallel to the anterior-posterior commissure plane. For all fMRI task participants, a high-resolution T1-weighted MPRAGE structural image was also obtained for atlas registration. For further details on acquisition parameters pertaining to each task, reference the original publication outlined in **Supp. Table 1**.

### fMRI data preprocessing

All fMRI studies included in the analysis implemented a uniform preprocessing stream, both for tasks and rest. More detailed overviews of processing steps can be found in the original publications corresponding to the tasks outlined in **Supp. Fig. 1**. Briefly, removal of noise and artifacts was performed using a series of automated steps. Temporal re-alignment was applied using sinc interpolation of all slices to the the first slice to account for differences in acquisition time of slices. Next, movement within and across BOLD runs was corrected using a rigid-body rotation and translation algorithm (Snyder 1996). To allow for comparison across subjects, whole-brain intensity normalization for each functional run was performed to achieve a modal value of 1000 across all voxels in the image (Ojemann et al. 1997). Functional data were resampled into 3 mm isotropic voxels and transformed into atlas space (Talairach and Tournoux 1988) in a single step. Using a series of affine transforms (Fox et al. 2005; Michelon et al. 2003), atlas registration was performed by aligning each subject’s T1-weighted image to a custom atlas-transformed (Lancaster et al. 1995) target T1-weighted template.

### Region and network definitions

In order to examine the functional role of the CO network, we first sought to identify regions with a high probability of association with this network in all subjects from each of our datasets. For that reason, we relied on the probabilistic mapping of functional brain networks as originally defined by Dworetsky et al. (2021); see probabilistic map of CO network in **Fig. 1B**). This approach identified 153 high-probability regions (see **Fig. 5A**), meaning these regions are assigned to a particular network in 75% of the large sample of young adults. Within these 153 regions, 15 regions were identified as belonging to the CO network with high probability (>75% consensus across individuals; mean [SD] peak probability across regions = 94.9% [4.6%]; see **Fig. 1C**).

### Task fMRI analysis: general linear model

A general linear model (GLM) was used to analyze activations in each of the fMRI tasks listed in **Supp. Table 1** (Miezin et al. 2000). In all task GLMs, a baseline constant and linear trend were included across each BOLD run to address drift effects. The mixed design tasks (Tasks 1–16) were analyzed with block-level regressors for sustained activity across the duration of each task, and events for onset and offset cues, correct trials, and error trials. Events were modeled with 8 timepoints using a finite impulse response approach (Ollinger et al. 2001a; Ollinger et al. 2001b). Activations for the blocked (boxcar) regressors were computed as the *z-*score relative to baseline; activations for events were computed as percent signal change relative to baseline.

The ‘slow reveal’ decision making tasks (Tasks 17–21; (Ploran et al. 2007; Ploran et al. 2011; Wheeler et al. 2008) were analyzed using an event-related approach, with separate event regressors for trials sorted by participant response time – i.e., regressors for participants responses at timepoints 4, 5, 6, and 7 (this selection captures the majority of trials; see (Gratton et al. 2017)). Each trial was again modeled using a finite impulse response approach with 16 timepoints. Activations for each event were computed and expressed as the percent signal change relative to baseline.

### Hierarchical clustering analysis of control-related task activations

A hierarchical clustering analysis (Cordes et al. 2002; Dosenbach et al. 2007; Salvador et al. 2005) was used to identify clusters of regions with similar task activation profiles related to task control. Seven different types of control-related task activations were included in this analysis (**Fig. 2**): (i) *<u>onset cues</u>* at the start of a block indicating the task to be performed (e.g.: “rhyme/no rhyme” in **Fig. 2B**), (ii) *<u>offset cues</u>*, indicating the end of the task set period (e.g., a red fixation cross in **Fig. 2B**), (iii) *<u>sustained signals</u>* elevated through an entire task block (green boxcar in **Fig. 2B**), (iv) *<u>errors</u>* committed by participants (e.g., responding “rhyme” to the word pair “fare/bank”; light gray signals in **Fig. 2B**), (v) *<u>ambiguity</u>* in a stimulus (e.g., a rhyme judgment about the word pair “bass/grace”; medium gray signals in **Fig. 2B**), relative to (vi) unambiguous *correct* trials (e.g., responding “no rhyme” to “fare/bank”) and finally (vii) *<u>decision-making</u>* timecourses (e.g., recognition judgment during a slow reveal task; yellow and orange signals in **Fig. 2C**; note that decision-making timecourses allow us to separate transient performance-related signals that may link to task parameter updates from gradual evidence accumulation responses (Gratton et al. 2017; Ploran et al. 2007; Ploran et al. 2011). These activations are related to the instantiation of task parameters (i,ii), maintenance of task signals (iii), and adjustment of signals based on ongoing performance (iv-vii).

For the analysis, timecourses (for, onset/offset cues, error/correct/ambiguous trials and slow reveal trials where participants indicated their responses as correct at steps 4, 5, 6, and 7) and z-scores of sustained task activation were extracted from each ROI, producing a total of 9 timecourses plus sustained activation per region from each task. For onset/offset-cue conditions and error/correct/ambiguous, activations came from two tasks (abstract/concrete, Task 1; and rhyme/non-rhyme, Task 2) which were modeled together to produce one timecourse per condition, each with 8 timepoints. Timecourses from the slow reveal task were extracted for each of 5 tasks (Tasks 17-21), each with 16 timepoints, and were then averaged across tasks. Sustained activations were modeled across a set of 16 different tasks. A list of the signals analyzed and information on their experiments’ parameters are shown in **Supp. Table 1** and **Supp. Table 2** respectively.

All timecourses and sustained activation values were concatenated for analysis. The following signals were included: (i-ii) onset/offset cue timecourses (8 timepoints per 2 signals; 8 × 2 = 16), (iii) sustained activation z-scores (sustained signals from 16 tasks), (iv-vi) error/ambiguous/correct timecourses (8 timepoints per 3 signals; 8 × 3 = 24), and (vii) slow reveal timecourses (16 timepoints per signal across 4 response bin timecourses; 16 × 4 = 64). We weighted these signals equally in analysis, resulting in a 1×768-datapoint vector per region (192 datapoints for each of i-ii, iii, iv–vi, and vii). A 15×768 matrix was then formed, containing a vector for each of 15 regions of interest (ROIs) from the CO network. A hierarchical clustering analysis was subsequently performed on this matrix, wherein the unweighted paired group method with arithmetic mean (UPGMA), included in the Statistics and Bioinformatics Toolbox in MATLAB (The MathWorks), was used to produce a dendrogram of region-wise relationships using the Euclidean distance between the concatenated vectors across regions. Using this same method, a supplementary clustering analysis was performed using data from the 15 CO ROIs along with 11 ROIs from the FP network and 3 from the Salience network. Additional analyses shown in the supplement plotted the consistency of these results across tasks.

### Resting-state fMRI analysis

To examine differences in the resting-state profiles of sub-networks produced by the clustering analysis, a Dartmouth dataset (see *Overview & Datasets*) was used. Preprocessing and functional connectivity processing steps for this dataset are described in greater detail by Gordon et al. (2016). Briefly, data underwent initial preprocessing to reduce artifacts as in the task fMRI data above, including slice-timing correction, whole-brain intensity normalization across each BOLD run to a modal value of 1000, motion correction via rigid body transformation, registration to a T1-weighted image and resampling to stereotactic atlas space (Talairach and Tournoux 1988) and isotropic 3mm voxels. Additional processing was performed to further denoise the data for functional connectivity analysis: data were demeaned and detrended, and nuisance signals (global signal, cerebrospinal fluid, white matter, six rigid-body motion regressors and their expansion terms) were regressed. High-motion frames (framewise displacement > 0.2 mm., along with short segments and segments at the start of each run) were interpolated over and a bandpass filter (0.08 – 0.009 Hz) was applied before spatially smoothing the data at FWHM (6mm).

The 69 subjects in this resting-state dataset were used to produce an average resting-state seedmap for each resultant cluster from the hierarchical clustering analysis. First, a group-average correlation matrix was produced by averaging all subjects’ voxel-to-voxel correlation matrices. From this group matrix, a seedmap for each region of interest was produced by averaging across voxels within the ROI, and then averaging across seedmaps of regions within a cluster. Voxels within 20mm of a given region were excluded from calculation of the mean to reduce the dependence on local signals strongly influenced by spatial auto-correlation. Cluster average seedmaps were qualitatively compared, and a difference seedmap was produced to contrast seedmaps from CO sub-network clusters.

A similar approach was also used to create a group average correlation matrix from the larger set of 153 high-probability ROIs. This was used to create a spring-embedding plot in order to visualize the community network structure of regions spanning the cortex, and particularly of our systems and sub-systems of interest. For this plot, the connection matrix was thresholded at a given edge density (i.e., top X% of correlation values). These were then subjected to a spring-embedding algorithm that places regions with more connections closer together in space, treating these connections as “springs” that are under tension (Kamada and Kawai 1989). For this analysis, the region-to-region matrix was thresholded at an edge density of 10% for the primary analyses, with supplemental analyses implementing thresholds at 8 and 12%.

In addition, the role of regions within each sub-network was quantified through the participation coefficient and within module degree metrics (Guimera et al. 2005). The participation coefficient was calculated for each node in the CO sub-systems as a means of measuring the degree to which a given region is “hub”-like, or well-connected to multiple networks. The participation coefficient is a measure of the diversity of a node’s connections across different networks, calculated for node *i* as:

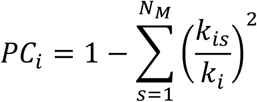

where *k*_*i*_ represents the node’s degree (number of connections to other nodes), *k*_*is*_ quantifies the node’s connections to other nodes within the same module *s*, and *N*_*M*_ represents the total number of modules in the graph.

The participation coefficient was calculated for each node correlation matrix across a range of thresholds (1% to 25%). Across the ROIs clustering into each sub-system, the mean participation coefficient was computed.

In addition, we sought to explore and compare the interconnectedness of control sub-system nodes to other nodes within their home network. This metric of within-module degree was computed as a z-score for each region; these calculations were also implemented across a range of 1% to 25% edge density thresholds and were again averaged across ROIs within an identified sub-system.

### Neurosynth meta-analysis

Finally, we identified common task-based associations from the literature for regions within each sub-system using the Neurosynth database (Yarkoni et al. 2011). Using its “associations” function, we entered the MNI-transformed coordinates of each sub-system’s ROI center coordinates and noted the resultant word associations drawn from meta-analysis maps (associations with the highest significant z-score were retained) that were commonly associated with that voxel. We manually excluded anatomical terms from this analysis, as the objective was to explore differentiation of function (not location) within sub-systems. Plural and singular terms were merged (e.g., ‘task’ and ‘tasks’ became ‘task(s)’). The word terms were used to produce a word cloud as a means of visualizing the groups of terms that are most commonly/repeatedly associated with regions within a given sub-system.

## RESULTS

### Overview

Our goal in this work was to provide a more detailed examination of the cingulo-opercular (CO) network and its role in task control by both (a) using an extended set of CO network regions with high consistency across individuals to increase the precision of CO definitions and (b) integrating information from a broad set of task activation and resting-state functional connectivity data. We begin by examining the consistency and diversity of CO region responses across a large set of task control signals using hierarchical clustering. These findings provide evidence for two major sub-system divisions within the CO network, only one of which is strongly associated with task control signals. We further interrogate this sub-system structure through an examination of meta-analytic task fMRI associations and functional connectivity of these regions. We close by contrasting task activation and functional network patterns of CO sub-systems with other higher-level ‘control’ systems of the brain.

### CO regions cluster into two sub-systems with apparently divergent roles in task control

We started by re-examining a large set of fMRI activations associated with task control (**Fig. 2**), in regions with high probability of being part of the CO network across people (**Fig. 1C**). We define task control signals as those associated with different aspects of instantiation, maintenance, and adjustment of task sets needed to achieve a goal (Logan and Gordon 2001).

In this work, we define task set initiation signals as (i) *<u>onset cues</u>* at the start of a block indicating the task to be performed (e.g., “abstract/concrete”) and (ii) *<u>offset cues</u>*, indicating the end of the task set period. We define task set maintenance as related to (iii) *<u>sustained signals</u>* elevated through an entire task block. Finally, we examine a range of performance-related signals that are indicators of a need for task set adjustments, including (iv) *<u>errors</u>* committed by participants (e.g., responding “abstract” to the word “cat”) and (v) *<u>ambiguity</u>* in a stimulus (e.g., “bass/grace” rhyming), relative to (vi) *<u>correct</u>* unambiguous trials (responding “concrete” to “cat”), as well as timecourses during (vii) *<u>decision-making</u>* trials (for further explanation, see *Methods* and (Gratton et al. 2017)).

We examined responses to all 7 signal types for each of the 15 high probability CO regions across 421 people in 21 tasks (**Supp. Table 1**). To identify common clusters of response types, we conducted a hierarchical clustering analysis (**Fig. 3**; see *Methods*). Two primary sub-systems emerged. The first sub-system (CO1, dark purple in **Fig. 3**) was composed of regions in the more rostral anterior insula (below the pars triangularis) and dACC/mFC. These regions had responses consistent with our original conceptualization of the CO network. As shown in **Fig. 3C-H**, these regions had strong responses to (i) onset and (ii) offset cues at the start and end of each block, elevated (iii) sustained signals across task blocks (aside from those that are perceptually driven as expected based on (Dubis et al. 2016), gray shading on right side of **Fig. 3E**), and higher responses to (iv) errors and (v) ambiguous stimuli relative to (vi) unambiguous correct trials. In addition, these regions showed (vii) large, relatively transient, response activations linked to the moment of decision during decision-making tasks (Gratton et al. 2017). These characteristics are consistent with these regions having a role in initiating (cues), maintaining (sustained), and adjusting (errors, ambiguity, transient decision response) task set signals, suggesting a broader role in task control.

**Figure 3:**
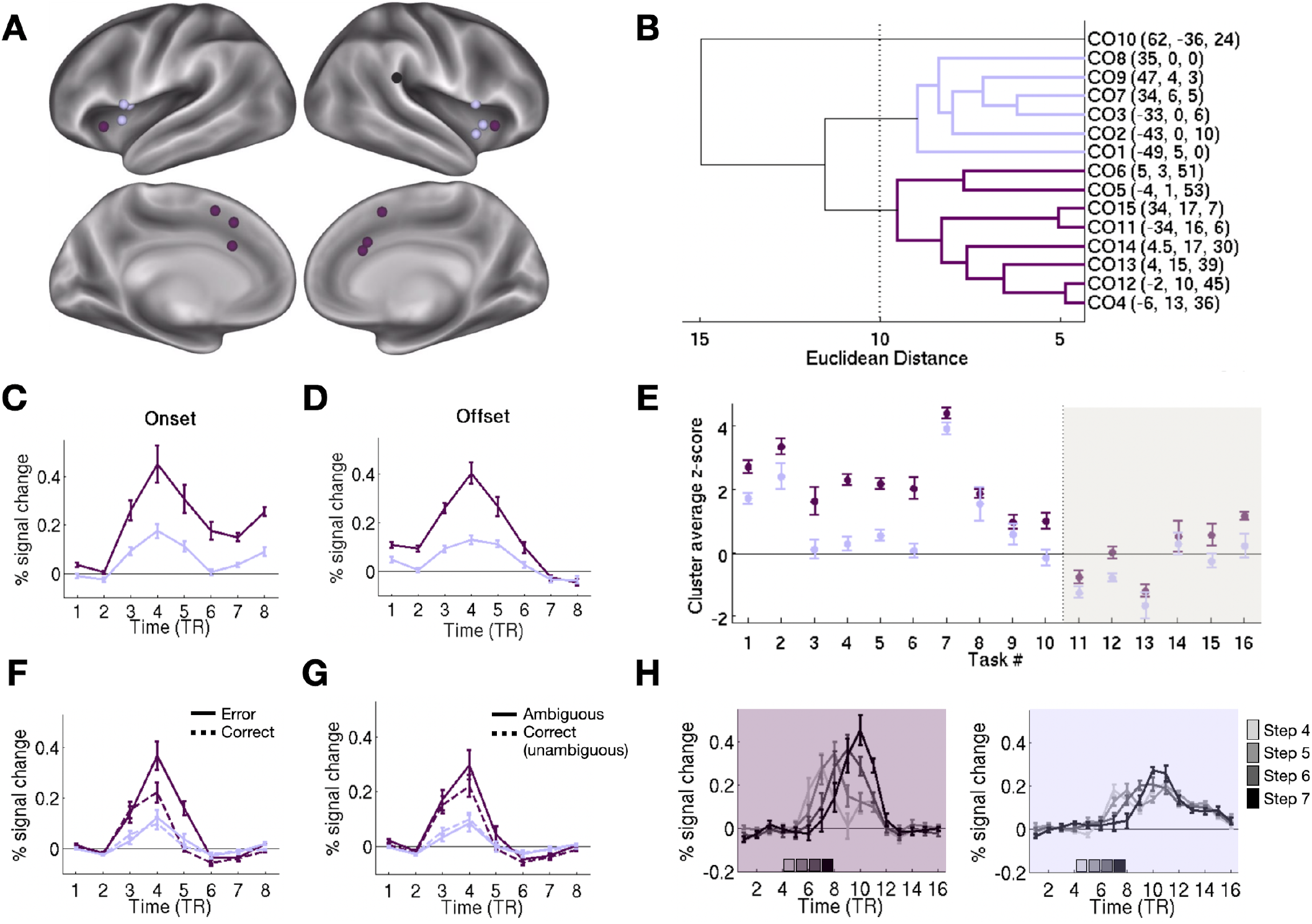
Distinct responses to control signals in CO network regions. We examined the profile of task control signals in 15 high probability CO network regions. Based on these responses, regions clustered into two distinct profiles: CO1 sub-system, shown in dark purple, and CO2 sub-system, shown in light purple (**A**: region locations; **B**: hierarchical clustering based on region responses). Responses are shown for the two sub-networks for signals related to task set initiation, such as **(C)** onset and **(D)** offset cues, **(E)** sustained signals related to task set maintenance, and task set adjustment signals such as responses to **(F)** errors vs. correct responses, **(G)** ambiguous vs. unambiguous correct responses, and **(H)** decision-making responses to a slowly revealed stimulus (steps 4-7 indicate the moment in the graded reveal at which the participant was able to identify a stimulus). Given the meta-analytic nature of the analyses, all error bars represent standard error of the mean across regions within each sub-system. Response characteristics separated by task are shown in ***Supp. Figs. 2*** and ***3***.

In contrast, the second sub-system (CO2, light purple in **Fig. 3**) had a fairly distinct profile. CO2 was composed of regions in a more posterior zone of the anterior insula, below and extending to the pars opercularis. These regions had relatively weak responses to (i) onset and (ii) offset cues (i.e., substantially smaller magnitude than those seen for CO1). (iii) Sustained response magnitudes were relatively low across most tasks, in all cases lower than that seen for CO1.

Unambiguous correct trials (vi) elicited weak positive responses, which were no larger for (iv) errors or (v) ambiguous stimuli. Finally, (vii) decision making responses in this region were low in magnitude and extended in time relative to the strong, transient responses at the moment of decision seen with CO1. Thus, the CO2 sub-system does not have signals strongly associated with task control (see also responses in CO1 and CO2 relative to other networks in **Supp. Fig. 1**; discussed further below). These differences between CO1 and CO2 responses were consistent across individual tasks (**Supp. Fig. 2-3**). A single additional CO region along the right supramarginal gyrus (black in **Fig. 3A**) showed a third profile that did not cluster closely with either CO1 or CO2 and did not show evidence of task control signals; see **Supp. Fig. 4** for closer characterization.

### CO sub-systems show diverging functional connectivity profiles

Next, we examined how the CO1 and CO2 sub-systems compared in their resting-state functional connectivity in a large sample of healthy young adults (Dartmouth 69; Gordon et al. 2016). A major preliminary piece of evidence that the CO network represents a distinct entity came from resting-state data (Dosenbach et al. 2007), which demonstrated that regions of the CO network had strong connectivity with one another, separate from regions in other networks (e.g., frontoparietal). Here, we see the same overarching pattern (**Fig. 4**): CO regions in both sub-systems show a relatively similar connectivity profile, with high correlations along the anterior insula and supramarginal gyrus, dACC/msFC, and anterior prefrontal cortex. Low correlations are seen with regions of the default mode network (posterior cingulate, angular gyrus), and intermediate correlations are seen with dorsal attention, somatomotor (e.g., highest in premotor areas), and visual regions. Functional connectivity for both CO1 (**Fig. 4A**) and CO2 sub-systems (**Fig. 4B**) look clearly distinct from those of other control-related systems like the frontoparietal and salience networks (**Fig. 4D and 4E**, respectively).

**Figure 4:**
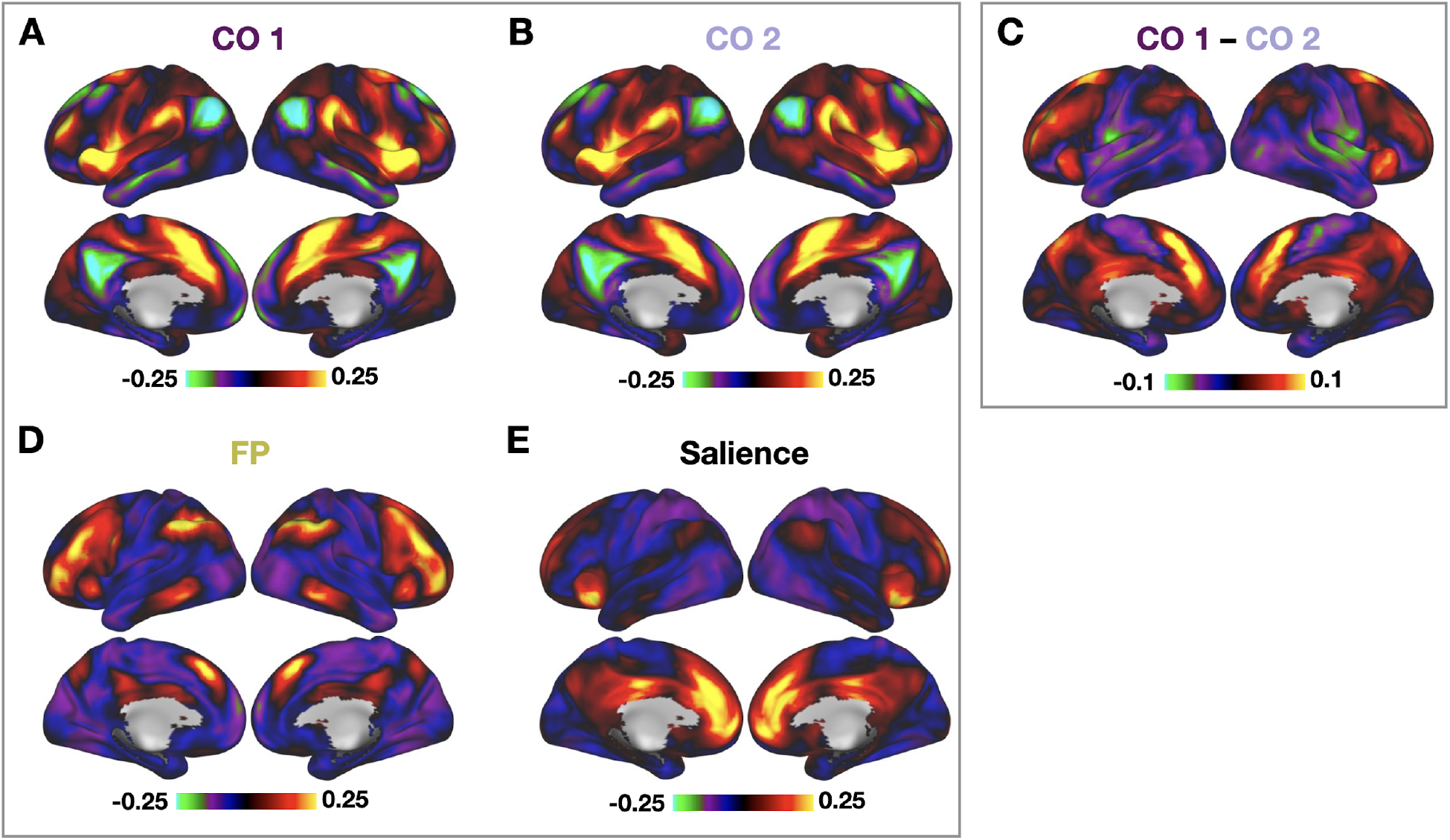
Resting-state functional connectivity of the CO1 **(A)** and CO2 **(B)** sub-systems. Both sub-systems had generally similar connectivity, consistent with their placement in the CO network. However, direct contrast between the two systems **(C)** reveals subtle distinctions in their functional network profiles. CO1 has higher connectivity to dACC/mFC, anterior and superior prefrontal, parietomedial, and dorsal attention locations, while CO2 has higher connectivity to mid and posterior insula, superior temporal, somatomotor, and extrastriate visual locations. Both CO sub-systems appear clearly distinct from the frontoparietal **(D)** and salience **(E)** networks.

Despite their overarching similarity, direct comparisons between the CO1 and CO2 systems revealed nuanced differences (**Fig. 4C**). Qualitatively, the CO1 sub-system has functional connectivity more akin to the “core” described earlier, with relatively higher functional connectivity with the dACC/msFC, anterior and superior frontal cortex and anterior insula within traditional CO regions. In addition, this region had relatively higher functional connectivity to regions of the parietomedial and dorsal attention networks. The CO2 sub-system had relatively higher connectivity to regions of the middle and posterior insula and superior temporal gyrus, and somewhat more elevated connectivity to primary somatomotor and extrastriate visual regions.

We next asked how these two sub-systems were positioned within the context of the full set of large-scale cortical networks. Resting-state functional connectivity was estimated among 153 high-consensus regions spanning 12 networks (Dworetsky et al. 2021). This functional connectivity matrix was thresholded (top 10% of functional connectivity edges; see other thresholds in **Supp. Fig. 5**) and used to create a spring-embedded layout, where nodes with a high number of connections are clustered relatively close to one another. As expected, both CO sub-systems (dark and light purple) clustered with one another and separate from high probability regions from other networks (**Fig. 5A**). The CO1 sub-system had connections interfacing with other higher-level systems like the frontoparietal (yellow), salience (black), and dorsal attention (green) systems, along with dense connectivity with its own network. CO2 regions appeared to have relatively fewer connections with other networks apart from the CO, suggesting it may have a more specialized role.

**Figure 5:**
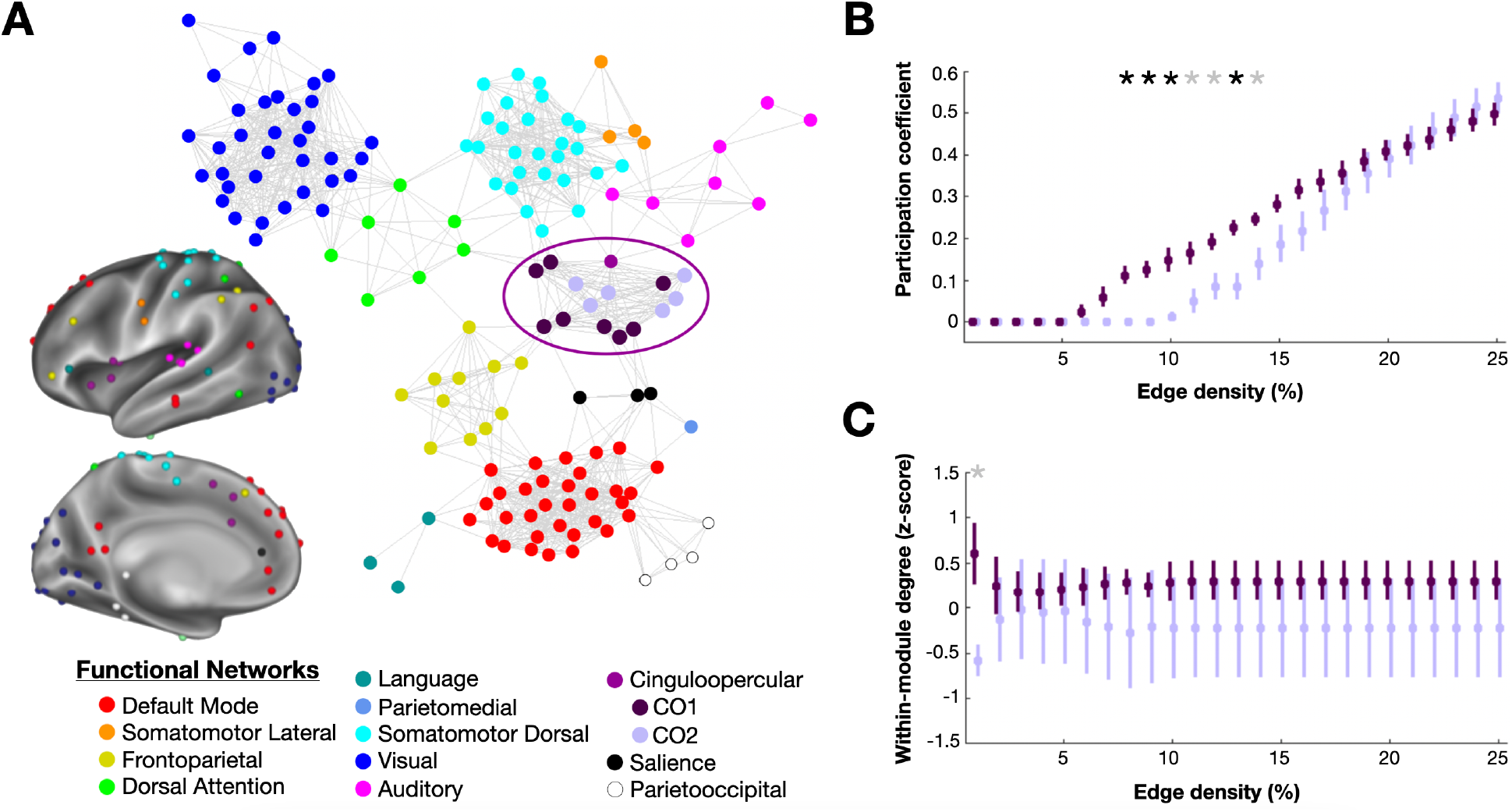
Functional connectivity of the CO sub-systems in the context of the whole brain ‘connectome’. **(A)** Functional connectivity was estimated among 153 regions (nodes) in 12 networks (colors; see inset for anatomical locations), generating a correlation matrix. This matrix was thresholded to include only the top 10% of connections (edges) and used to generate a spring embedded graph (see *Methods*; see graphs at other thresholds in **Supp. Fig. 5**), where more highly interconnected nodes are placed closer to one another. CO network regions (circled) are colored in shades of purple (CO1 = dark purple; CO2 = light purple, posterior CO node = magenta). CO regions all clustered with one another, with CO1 regions appearing to show high connectivity to regions both in the CO cluster and in other networks (frontoparietal, dorsal attention, salience); CO2 regions appeared relatively more segregated. The single CO region from the posterior insula (magenta) was most closely aligned with CO2 regions and the auditory network. **(B)** The participation coefficient, a measure of whether a node has connections to multiple different networks, was used to quantify the extent to which CO1 and CO2 regions served as connectors across a range of thresholds for the graph (edge density). CO1 regions (dark purple) had relatively higher participation coefficient values than CO2 regions, suggesting that they are stronger connectors (indicated as a black star = p<0.05 after FDR correction for multiple comparisons; a gray star indicates differences that did not achieve significance after FDR correction). **(C)** Within-module degree, a measure of a region’s connectivity to other same-network nodes, was used to estimate whether CO1 or CO2 regions serve as stronger within-network hubs. Within-module degree does not appear to differ strongly between CO1 and CO2 regions.

To quantify these apparent differences in connectivity we used the graph theoretical properties of participation coefficient and within-module degree. The participation coefficient is a metric that quantifies the extent to which a region has high between-network connections to multiple diverse networks (acting as a ‘connector’), whereas within-module degree quantifies the extent to which a region has many within-network connections (acting as a ‘module hub’). For participation coefficient, values closer to 1 indicate that a region has connections distributed across multiple networks, values closer to 0 indicate that a region has connections concentrated within a single network (Guimera et al. 2005). Past work has suggested that the CO network is relatively enriched in connectors (Gratton et al. 2018b; Power et al. 2013), a quality we hypothesize is important for instantiating diverse parameters during task control (Gratton et al. 2018b). Here we show that the participation coefficient of regions in the CO1 sub-system (dark purple) is higher than that for regions in the CO2 sub-system (light purple) for a range of graph thresholds (**Fig. 5B**; thresholds exhibiting a significant difference [adjusted p<0.05 obtained from two-tailed t-tests and after FDR correction for multiple comparisons] between clusters are denoted with a black asterisk; thresholds marked with a gray asterisk indicate differences not passing FDR correction). This suggests that the CO1 sub-system is composed of stronger between-network connectors than the CO2 sub-system, consistent with a role in coordinating task control information across systems.

One question is whether the CO1 sub-system is more external to the network, carrying inter-network connections, while the CO2 sub-system is more central to integrating information within its own network. Within-module degree is a normalized measure (z-score) of the number of connections a region has to other regions within its own network (Guimera et al. 2005); values over 0 indicate more than the average number of intra-network connections for the network. The within-module degree, therefore, measures the extent to which a region can be considered a within-network or ‘module hub’. This measure may help identify regions especially important for integration of information within a single system. Contrary to the initial hypothesis, the two CO sub-systems don’t differ strongly on their within-network connections, as demonstrated by their similar within-module degree (**Fig. 4C**); in fact, the CO1 system has numerically higher values, especially at sparse thresholds. The significantly higher participation coefficient and numerically higher within-module degree suggests that the CO1 has a stronger integrator role, especially for coordinating between networks, but also potentially integrating this information within the CO network; the CO2 sub-system, in contrast, is relatively more isolated, perhaps in association with a more specialized functional role.

### An exploratory meta-analysis of the CO sub-system suggests divergent functions

The previous analyses confirmed that the CO1 sub-system has a task response and functional connectivity profile consistent with task control: these regions exhibit a range of task control signals across multiple different tasks along with connector-like functional connectivity to multiple networks. However, these data also revealed the distinct CO2 sub-system, with much weaker representation of control signals and subtly divergent functional connectivity patterns. These results beg the question: to what extent do the CO1 and CO2 distinctions replicate in the broader literature, and can this provide hints to the underlying function of the CO2 sub-system?

To address this question, we conducted an exploratory analysis of the CO1 and CO2 regions using Neurosynth to identify common functional terms associated with each region. Coordinates from each region in MNI space were entered into Neurosynth’s location search feature. Resultant keywords (word associations with a significant z-score) were then entered into a word cloud for all the regions within a sub-system, with larger sizes indicating key words that occurred more frequently (note singular/plural terms, e.g., ‘task’ and ‘tasks’, were grouped; anatomical terms were removed, see *Methods;* see **Supp. Table 3 and 4** for a full list of *Neurosynth* associations, including anatomical terms, for regions in CO1 and CO2 respectively). **Figure 6** shows the result of this analysis separately for each sub-system. While some terms overlapped, the CO1 sub-system had a higher frequency of abstract terms related to control (e.g., “task(s)”, “working memory”, “demands”, “load”, “conflict”), including less common terms (e.g., “stroop”, “sustained attention”, “effortful”, “preparation/preparatory”, “inhibition”) that were not evident at all in CO2. The CO2 system did not show a high frequency of these abstract task control terms, being instead dominated by phrases related to motor function (e.g., “movement(s)”, “finger”, “force”, “motor imagery”,) and pain/interoception (e.g., “pain”, “painful”, “noxious”, “autonomic”). Some of these motor and pain related terms were present in the CO1 sub-system as well, but with less frequency (i.e., these are not in the top 5 for the CO1 sub-system as they are for CO2). These qualitative results provide evidence for distinctions between the CO1 and CO2 sub-systems from the broader literature, including further support for a role of the CO1 sub-system in task control, and preliminary evidence that the CO2 sub-system may have a role in pain/interoception and motor functions.

**Figure 6:**
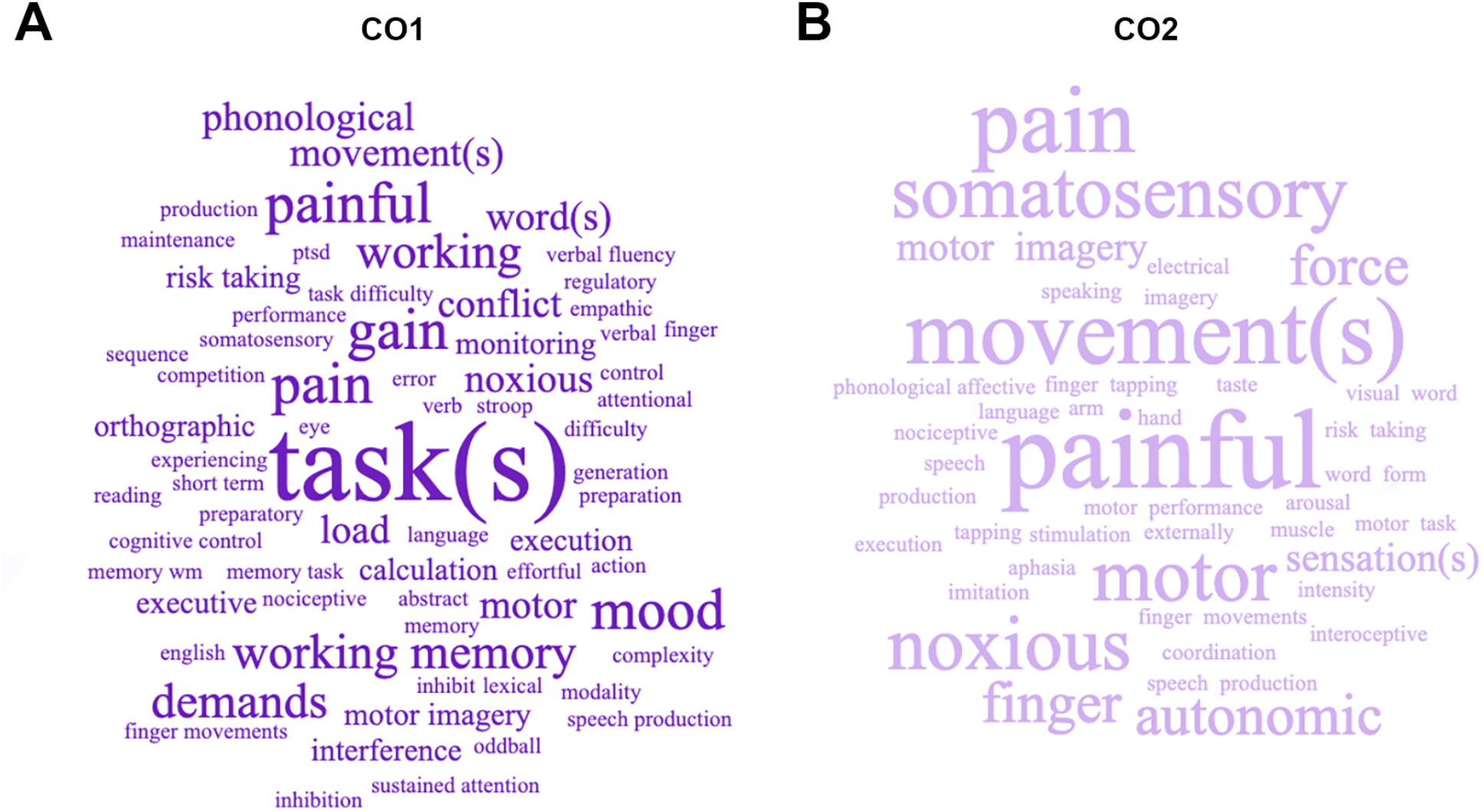
Exploratory analysis of functional terms associated with (A) CO1 and (B) CO2 regions. Neurosynth was used to identify keywords associated with the regions in each sub-system, pictured here as a word cloud, with keyword size proportional to its frequency. While some common terms are seen, the two sub-systems are dominated by different categories of items: CO1 has more terms associated with task control, while CO2 has more terms associated with pain/interoception and motor function.

### Relationship between CO sub-systems and other ‘control’ networks

Thus far, we have focused on common and distinct signals within the CO network and shown evidence for two differentiable CO sub-systems. We next asked how these attributes compare with other putative ‘control’ networks, specifically the frontoparietal and salience networks that have been most often paired with the CO (Dosenbach et al. 2007; Gratton et al. 2018b; Menon and Uddin 2010; Seeley et al. 2007; Uddin et al. 2019); note that here we are enumerating each system based on our own network nomenclature, which may show differences with nomenclature across other groups (Uddin et al. 2019). Given their proximity, and variability in network distributions across individuals (**Fig. 1**; (Dworetsky et al. 2021)), we once again focus only on regions with a high consistency of assignment to each functional system across people.

Resting-state functional connectivity analyses show distinct spatial topographies to CO, frontoparietal, and salience network regions (**Fig. 4**). The differences in functional connectivity between these three networks are substantially larger than those seen between the CO1 and CO2 sub-systems. This is further borne out in the graph layouts of the large-scale brain networks (**Fig. 5A**), where CO1 and CO2 sub-systems cluster together, nearby but separate from frontoparietal and salience regions.

Task responses across the three control systems show relatively more subtle distinctions compared to resting-state functional connectivity (**Supp. Fig. 1**). As a whole, FP regions show similar magnitude (i) onset and (ii) offset signals as the CO2 regions, smaller than CO1. (iii) Sustained signals are of intermediate magnitude. (vi) Correct trial signals are small, but larger for (iv) errors and (v) ambiguous stimuli, as in CO1. (vii) Decision-making responses have early and graded responses, that peak around the moment of decision, consistent with a role in evidence accumulation (Gratton et al. 2017), which is categorically different from CO1 and CO2. Salience regions also have small (i) onset and (ii) offset cue signals that are similar to CO2, (iii) Sustained signals were variable across tasks, but mostly smaller than was evident for CO1. There was no clear response to (vi) correct or (v) ambiguous trials, but a slightly elevated response to (iv) errors. (vii) Decision-making responses were small and delayed, unlike CO and FP profiles. These differences/similarities between the control systems were consistent across individual tasks (**Supp. Figs. 2-3**; see also **Supp. Fig 6** for the hierarchical analysis repeated for regions from all 3 networks). Thus, functional responses in task control show differences across the three control systems, but with more subtle distinctions relative to those seen in the resting-state functional connectivity data. These results emphasize the value of uniting both task and resting state fMRI data in the analysis of large-scale brain systems.

## DISCUSSION

In our previous work, we have proposed that the cingulo-opercular (CO) network is central to task control (Dosenbach et al. 2006; Gratton et al. 2017; Gratton et al. 2018b; Neta et al. 2017; Neta et al. 2014; Power and Petersen 2013). However, many alternative explanations have also been provided for the functions of the CO network from a wide variety of domains and disciplines (Alexander and Brown 2011; Bartels and Zeki 2004; Brown and Braver 2007; Craig 2009; Grinband et al. 2011; Lieberman and Eisenberger 2015; Neta et al. 2013; Sadaghiani and D’Esposito 2015; Sterzer et al. 2002; Thompson-Schill et al. 1997; Wessel et al. 2012). One putative explanation for this variability, largely overlooked until recently, is that the differing explanations arise from heterogeneity in brain networks (Finn et al. 2015; Mueller et al. 2013; Seitzman et al. 2019; Smith et al. 2021). In this work, we accounted for inter-subject variability by leveraging our newly minted regions that have a high probability of CO network assignment across people. We explored the consistency of our original characterization of the CO network in supporting task control through an examination of a variety of task signals related to the initiation, maintenance, and adjustment of task demands, as well as resting-state correlations, and a meta-analysis of task reports. Together, these findings provide evidence for two distinct CO sub-systems, only one of which is strongly associated with task control as we previously conceived it.

### Divergence in the CO network: evidence for two sub-systems

The current work provides evidence of two related but functionally divergent sub-systems within the CO network: CO1 and CO2 (**Fig. 3**). Both CO1 and CO2 have similar functional connectivity profiles, appearing as part of the same network and distinct from other systems, consistent with past descriptions (Dosenbach et al. 2006; Power et al. 2011; Yeo et al. 2011); **Fig. 4**). However, task control responses show clear distinctions between the CO1 and CO2 and these are accompanied by more subtle differences in network architecture. These results fit with prior evidence of a diversity of signals in nearby regions of the CO network (Klugah-Brown et al. 2022; Nee et al. 2007; Nelson et al. 2010; Smith et al. 2021) and may provide insight into the many alternate views on the role of the CO/dACC.

For example, in social neuroscience, some have argued that the dACC is important for social pain or rejection (Lieberman et al. 2016; Lieberman and Eisenberger 2015); but see (Wager et al. 2016). In other cross-disciplinary work, it has been argued that the dACC is important for social, cognitive, and affective conflicts (Botvinick et al. 1999; Botvinick et al. 2001; Carter et al. 1998; Cassidy et al. 2017; Hehman et al. 2014; MacLeod and MacDonald 2000; Milham et al. 2001; Ochsner et al. 2009; Stolier and Freeman 2017), or decision uncertainty or ambiguity (Neta et al. 2013; Sterzer et al. 2002; Thompson-Schill et al. 1997). In particular, the dACC has been associated with novelty (Wessel et al. 2012), prediction of response outcome (Alexander and Brown 2011) risk (Brown and Braver 2007), and even simply increased reaction time (Grinband et al. 2011). Meanwhile, the aI/fO has been associated with empathy (Singer et al., 2004; Singer, 2006), interoception, and affiliation (Bartels and Zeki 2004), and has been proposed as playing a central role in a model of autism (Uddin and Menon 2009). Not surprisingly, then, these regions have been the victim of controversial reverse inferencing. Here we demonstrate that there may be a division between CO regions, with only a subset having strong involvement in task control functions.

Past work has also suggested that diverse functional accounts of the CO may arise from subdivisions within these regions of the brain. For example, one prominent early suggestion was that the anterior cingulate is subdivided into separate regions for emotional/pain and cognitive regulation (Bush et al. 2000), although later investigations suggested significant overlap in these signals in the dACC/mFC (Craig 2009; Luu and Posner 2003; Medford and Critchley 2010; Shackman et al. 2011), and even that systems for emotion and cognition are not distinguishable (Pessoa 2008). Differences in specific forms of control processes (e.g., related to conflicts, errors, switching) have also been linked to overlapping but dissociable sub-regions of the dACC/mFC (Nee et al. 2011). Work from Nelson et al. (2010) reported distinctions in task activations and connectivity of sub-regions of the anterior insula (see also recent work by (Klugah-Brown et al. 2022)). However, it is unclear if divisions in this prior work reflected divisions among large-scale systems (e.g., between the CO and salience along the dorsal/ventral anterior insula, CO and frontoparietal along dorsal and ventral regions of the dACC/mFC) rather than subdivisions within the same functional system, and to what extent they might be driven by variation in functional network locations across people. Here, we demonstrate that, after addressing individual variation in network locations, the CO shows clear divisions in its task control responses, especially along the anterior insula; these are of a different flavor than the distinction seen across functional systems (e.g., CO vs. salience, which show much more pronounced functional connectivity differences, and a different form of task control responses **Fig. 4, Supp. Fig. 1**; discussed more in detail below).

#### CO1: A task-control sub-system

In this work, the CO1 system appeared to act as the ‘connector and integrator’ of the CO network, with a strong role in task control. This sub-system was composed of regions in the more rostral anterior insula and the dACC/mFC. During tasks, this sub-system had a number of responses consistent with a role in initiating, maintaining, and adjusting task parameter signals (Logan and Gordon 2001), including: strong responses to onset and offset cues of tasks, sustained signals during task periods, and responses to errors, ambiguity, and decision points (**Fig. 3**). These responses match our original characterization of the CO ‘task control’ system (Dosenbach et al. 2006; Gratton et al. 2017; Neta et al. 2014), and also generally fit with prior models of the importance of the dACC/mFC in control-related functions (e.g., (Botvinick et al. 2001; Jonides and Nee 2006; Miller and Cohen 2001)). Indeed, Neurosynth guided meta-analysis found a high proportion of terms associated with task control and executive functions for CO1 regions (**Fig. 6**).

In addition to its task responses, CO1 regions had a relatively high degree of connectivity to other networks during rest (while maintaining strong within-network connectivity). This architecture indicates that CO1 regions are well positioned to act as connectors and integrators among diverse brain networks. Past work has theorized that a connector-like role is critical for task control, which requires a need to integrate information across regions specialized for different functions in a flexible manner depending on task demands (Bertolero et al. 2015; Bertolero et al. 2017; Gratton et al. 2018b). The high connector status of the CO1 sub-system maps onto the findings of Power and colleagues (2013) which suggested that the CO network is enriched in connector hubs of the brain. Altogether, the CO1 sub-system has properties largely consistent with previous descriptions of the CO network and its role in task control. However, previous studies, both of task activation and connectors, did not account for the variability and extent in CO network regions across people. By addressing these factors in this work (through the selection of high-consensus CO locations), we are able to demonstrate that these previous properties are relatively specific to only one subset of the CO network.

#### CO2: A sub-system with weak control, but preserved motor and pain responses

The CO2 sub-system diverged functionally from the CO1 and these prior descriptions of the CO ‘control’ network, with substantially less evidence for a role in task control. While still clearly part of the CO network based on functional connectivity, the CO2, which is composed of regions in the more caudal anterior insula, had relatively weak signals related to task initiation, maintenance, and adjustment (**Fig. 3**). While showing similar within-network connectivity as the CO1, the CO2 sub-system had relatively weak connectivity with other networks (**Fig. 5**), suggesting that it may have a more isolated and specialized role. Meta-analytic results (**Fig. 6**) suggest that the CO2 sub-system is most closely related to motor, pain, and interoceptive functions, and thus might serve as a form of internal regulation that supports task control.

These findings fit with past work suggesting that regions in the CO may be integral to motor control (Grinband et al. 2011; Newbold et al. 2021) and internal regulation (Craig 2009; Lieberman and Eisenberger 2015; Luu and Posner 2003; Medford and Critchley 2010; Mesulam and Mufson 1982). The insula has been hypothesized to have a posterior to anterior functional organization with the posterior insula responsible for representations of the affective feelings from the body, the mid-insula integrating these representations with inputs from other sources, and the anterior insula producing a moving window of subjective feelings. Homeostatic feelings instantiated in the more posteriorly oriented CO2 might inform action selection and online adjustments of control carried out by CO1. In addition, recent work has suggested that the CO network may play an important role in motor plasticity (Newbold et al. 2021), consistent with motor-related terms associated with CO2. However, given overlap in these meta-analytic terms across CO1 and CO2 regions (**Fig. 6**), these functions may be carried out in conjunction with CO1. Additional work will be needed to better understand the unique role of CO2 regions, with tasks probing and contrasting task control, motor, emotion, and interoceptive functions within these regions.

### What is the unifying link within the CO network?

Although the CO1 and CO2 sub-systems are dissociable, within the context of the whole brain, these sub-systems still share many features. Most dominantly, both fit cleanly within the CO network based on their functional connectivity patterns. Much like simpler and better studied neurobiological systems (visual, motor; (Biswal et al. 1995)), CO regions across both sub-systems have highly-coherent spontaneous activity. These CO regions show much lower spontaneous correlations with regions of other association systems, including the frontoparietal and salience networks. Thus, the CO acts as a coherent and distinct resting-state network. As part of the same network, we argue that the two sub-systems likely have related functional properties (like visual and motor systems), linked to genetic/developmental constraints (e.g., (DiNicola and Buckner 2021)) and a history of co-activation (Laumann and Snyder 2021; Petersen and Sporns 2015). But, an open question is what common property unites and activates these disparate CO sub-systems.

In our detailed exploration of various task control signals, we were unable to reveal a profile that provided a particularly strong unifying account for all CO region functions that was distinct from functions associated with other high-level networks. One possibility is that all CO regions will show common responses to other signals related to control, such as working memory, response inhibition, and visual attention that were not studied in detail in this manuscript. Having said that, our literature-based meta-analysis using the Neurosynth database found a relative enrichment in CO1 for abstract control-related terms, including many of these alternative processes that were incorporated in the tasks we included, suggesting that this possibility is not well supported by our current evidence.

Although speculative, another possibility is that the sub-system structure of the CO provides hints as to its unifying components, linked to progressive levels of abstraction in control. We hypothesize that the CO system may have originated as a system specialized for relatively basic body-level control related to interoception, pain, and motor actions (as discussed in the previous section). While this may remain a specialized component of CO functions (carried out by the CO2 sub-system), some regions of the CO network (CO1 in our proposal) may have evolved to additionally represent abstract task parameters and carry out domain-general aspects of control. This proposal is in analogy to theories regarding the evolution of the frontal eye fields (and broader dorsal attention system) from motor regions involved in directing eye movements, toward a more abstract role in directing attention even in the absence of changes in eye position (Awh et al. 2006; Corbetta et al. 1998; Rizzolatti et al. 1987). If true, this speculation would suggest that tasks focused on body-level control would recruit both CO1 and CO2, while task control requiring integration or translation across a range of domains would be linked more exclusively to CO1, necessitating additional links to sensory, memory, and other domains. Future work will be needed to explicitly test this hypothesis.

### Distinctions among large-scale ‘control’ systems

Despite their differences, the CO sub-systems appear clearly distinct from other higher-level ‘control’ systems. Consistent with past work (Gordon et al. 2017; Power et al. 2011), resting-state correlation patterns were strongly and consistently different between CO, frontoparietal, and salience regions (**Fig. 4**). Task control responses were also differentiable as a whole (**Supp. Fig. 1**), but these differences were more subtle, and prone to exceptions for specific regions (**Supp. Fig. 6**; note that one frontoparietal and one salience region clustered with CO1 and CO2 respectively). Thus, although our original conception of functional differentiation between CO and frontoparietal networks was more or less all or none (Dosenbach et al. 2008; Dosenbach et al. 2007; Power and Petersen 2013), the more recent analyses make the distinction in task control signals more a matter of degree than categorical (see also (Assem et al. 2020)), and hint at the presence of sub-system structure within these networks as well.

Future work will be particularly needed to carefully disentangle the properties of the CO from the salience network, which are often conflated in the literature. For example, there is substantial overlap in the regions designated as parts of the CO and salience networks across papers (and still others grouping these and the frontoparietal network in a more general executive function network) (Gratton et al. 2018b; Uddin et al. 2019). Thus, the taxonomy used in a particular paper is not a guarantee of reference to the same (or distinct) functional/anatomical structure (Uddin et al. 2022). However, in our hands resting-state analyses consistently find strong distinctions between two nearby systems (Gordon et al. 2017; Power et al. 2011) that we refer to as salience and CO, especially in studies that respect individual variability in system locations (Gordon et al. 2017) (the salience system is smaller and more variable across people, leading to more difficulty in identifying it in group analyses; note the smaller number of consistent salience regions in our probabilistic atlas). Additional insights into the differences between these two systems will therefore require approaches that not only carefully address anatomical definitions, but also that account for individual differences in the locations of these functional networks.

### Limitations and Future Research Directions

The current manuscript builds on our past theories about the role of the CO network by carefully accounting for individual differences while investigating the functional and network properties of CO regions. In this enterprise, we discovered (somewhat to our surprise) the presence of a novel CO2 sub-system which didn’t align closely with our previous theoretical stance. As such, these findings are limited in certain dimensions which motivate future research directions.

First, in this work, we largely focused on task parameter signals associated with task control, based on our previous research/theoretical leanings. However, these task responses did not provide a clear picture of a functional role specific to CO2. Thus, our ideas for the role of CO2 are speculative thus far, driven by a terminology-based meta-analysis of these regions. Future work will be needed to test our speculative considerations about the potential role of these regions in motor, pain, and interoceptive functions. In addition, as noted in the previous section, it will likely be helpful to more fully characterize the space of control signals to better understand the sensitivity of CO1, CO2, frontoparietal, and salience regions to other variables of interest (working memory, response inhibition, etc.).

Second, while we examined regions with high consistency in network labels across people, our task analyses were limited to probabilistic statements about the network assignments of regions (e.g., looking at CO regions that are labeled as CO in many, but not all, individuals). Future work with networks defined in single individuals and contrasting signals within those individuals may help to disambiguate apparent overlap between regions. Furthermore, these analyses may allow us to probe anatomical regions that were excluded here due to their variable locations across individuals (e.g., the often found, but inconsistent anterior dorsolateral PFC region associated with the CO network).

## Conclusions

A substantial debate continues regarding the functional role of CO regions, particularly the dACC/msFC, and the bilateral aI/fO. We have previously proposed that these regions have a role in task control, while others have proposed a range of functions related to social, emotional, and cognitive domains. A lack of precision in the anatomical locations of the CO network – especially in light of other, closely juxtaposed networks, all of which vary spatially across people – suggests a potential cause for these differing accounts. Here, we use regions that have high correspondence across people to show that the CO network is composed of two divergent sub-systems. One has a role in task control, consistent with our previous models. The other does not. Both appear to be part of a single network at rest that is distinct from other putative ‘control’ networks, but the two sub-systems differ in their positions within the CO network structure. These results are supported by a broad set of fMRI task and rest data, along with meta-analytic results from the broader literature. Thus, these findings suggest that the CO network should be reconceptualized as a broader system with at least two sub-components, only one of which is closely tied to task control, the other of which may be tied more closely to body-oriented processing. This evidence leads us to propose novel theories about the heterogeneity and unity of the CO network.

## ACKNOWLEDGMENTS

This work was supported by funds from the NSF CAREER BCS-2048066 (CG), CAREER BCS-1752848 (MN), R01MH118370 (CG), R01MH111640 (MN), and the Therapeutic Cognitive Neuroscience Fund (DMS). We would also like to thank Dr. Derek Nee, Jessica Wood, and Dr. Regina Lapate for helpful conversations on this work.

## Footnotes

1 Importantly, many studies reporting activation in dACC show a larger swath of activity that extends dorsally into the medial superior frontal cortex (msFC) or pre-supplementary motor area (pre-SMA; Brown & Braver, 2005; O’Reilly et al. 2013). For example, Lieberman and Eisenberger (2015) used Neurosynth to demonstrate that search terms “dACC” and “anterior cingulate” resulted in activation maps that included this more dorsal region. Thus, as we discuss the functional role of the CO network, including dACC, we refer to this broader region of dACC and adjacent msFC.

## Notes

### Competing Interest Statement

The authors have declared no competing interest.

