## Supplementary Material for "The cingulo-opercular network is composed of two distinct sub-systems"

### SUPPLEMENTAL TABLES

**Supp. Table 1**: Description of all tasks used for analysis of fMRI task control signals. *Note: asterisks in the 'Task' column indicate pairs of studies using the same set of subjects. Pairs with the same subjects include tasks 1 and 2 (*); 3 and 15 (**); 4 and 16 (***); 11 and 12 (****).*

| # | Task description | Publication (original) | Task design | Stimuli | Input modality | Output modality | N | Signals used |
|---|---|---|---|---|---|---|---|---|
| 1 | Abstract/ concrete (Auditory, 3T)* | Neta et al., 2014 | Mixed block/ event-related | nouns | auditory | button | 34 | error/ambiguous/ correct timecourses; sustained Z-score; onset/offset cues |
| 2 | Rhyme/ non-rhyme* | Neta et al., 2014 | Mixed block/ event-related | words | visual | button | 34 | error/ambiguous/ correct timecourses; sustained Z-score; onset/offset cues |
| 3 | Abstract/ concrete (Auditory, 1.5T)** | Dosenbach et al., 2006 | Mixed block/ event-related | nouns | auditory | button | 24 | sustained Z-score |
| 4 | Abstract/ concrete (Visual)*** | Dosenbach et al., 2006 | Mixed block/ event-related | nouns | visual | button | 17 | sustained Z-score |
| 5 | Cross-modal attention | n/a | Mixed block/ event-related | words | auditory/ visual | button/ speech | 32 | sustained Z-score |
| 6 | Living/ Non-living | Dosenbach et al., 2006 | Mixed block/ event-related | images | visual | button | 34 | sustained Z-score |
| 7 | Motor timing | Dosenbach et al., 2006 | Mixed block/ event-related | tone patterns | auditory | button | 32 | sustained Z-score |
| 8 | Object naming | Dosenbach et al., 2006 | Mixed block/ event-related | images | visual | speech | 18 | sustained Z-score |
| 9 | Noun/ verb | Dubis et al., 2014 | Mixed block/ event-related | nouns/ verbs | visual | button | 30 | sustained Z-score |
| 10 | Sustained task load | n/a | Mixed block/ event-related | words/ images | auditory/ visual | button | 28 | sustained Z-score |
| 11 | Glass pattern (2-level)**** | Dubis et al., 2016 | Mixed block/ event-related | dot pairs | visual | button | 20 | sustained Z-score |
| 12 | Glass pattern (4-level)**** | Dubis et al., 2016 | Mixed block/ event-related | dot pairs | visual | button | 20 | sustained Z-score |
| 13 | Glass patterns (limited) | Dubis et al., 2016 | Mixed block/ event-related | dot pairs | visual | button | 30 | sustained Z-score |
| 14 | Visual attention | n/a | Mixed block/ event-related | Gabor patches | visual | button | 30 | sustained Z-score |
| 15 | Visual search** | Dosenbach et al., 2006 | Mixed block/ event-related | Gabor patches | visual | button | 24 | sustained Z-score |
| 16 | Upper/ Lower-case*** | Dosenbach et al., 2006 | Mixed block/ event-related | nouns | visual | button | 17 | sustained Z-score |
| 17 | Object identification | Ploran et al., 2007 | Event-related (slow reveal) | object images (gradual dissolve) | visual | button | 13 | slow reveal timecourses |
| 18 | Word identification | n/a | Event-related (slow reveal) | words (gradual letter reveal) | visual | button | 13 | slow reveal timecourses |

| | | | | | | | | |
|---|---|---|---|---|---|---|---|---|
| 19 | Object priming | n/a | Event-related (slow reveal) | object images (gradual dissolve) | visual | button | 24 | slow reveal timecourses |
| 20 | Object retrieval | n/a | Event-related (slow reveal) | object images (gradual dissolve) | visual | button | 26 | slow reveal timecourses |
| 21 | Object shuffle | Ploran et al., 2011 | Event-related (slow reveal) | object images (jittered mask) | visual | button | 16 | slow reveal timecourses |
| 22 | Resting state | Gordon et al., 2016 | n/a | n/a | n/a | n/a | 69 | functional connectivity |

**Supp. Table 2:** Description of MRI acquisition parameters for all tasks.

| # | Name | TR (s) | Voxel size | Scanner strength |
|---|---|---|---|---|
| 1 | Abstract/concrete (Aud/3T) | 2.5 | 4 x 4 x 4 | 3T |
| 2 | Rhyme/non-rhyme | 2.5 | 4 x 4 x 4 | 3T |
| 3 | Abstract/concrete (Aud/1.5T) | 2.5 | 3.75 x 3.75 x 8 | 1.5T |
| 4 | Abstract/concrete (Vis) | 2.5 | 3.75 x 3.75 x 8 | 1.5T |
| 5 | Cross-modal attention | 2.5 | 3.75 x 3.75 x 8 | 1.5T |
| 6 | Living/nonliving | 2.5 | 3.75 x 3.75 x 8 | 1.5T |
| 7 | Motor timing | 2.63 | 3.75 x 3.75 x 8 | 1.5T |
| 8 | Object naming | 3.18 | 3.75 x 3.75 x 8 | 1.5T |
| 9 | Noun/verb | 2.5 | 4 x 4 x 4 | 3T |
| 10 | Sustained task load | 2.5 | 4 x 4 x 4 | 3T |
| 11 | Glass pattern errors (2-level) | 2.5 | 4 x 4 x 4 | 3T |
| 12 | Glass pattern errors (4-level) | 2.5 | 4 x 4 x 4 | 3T |
| 13 | Glass patterns (limited) | 2.5 | 4 x 4 x 4 | 3T |
| 14 | Visual attention | 2.5 | 4 x 4 x 4 | 3T |
| 15 | Visual search | 2.5 | 3.75 x 3.75 x 8 | 1.5T |
| 16 | Upper/lowercase | 2.5 | 3.75 x 3.75 x 8 | 1.5T |
| 17 | Object identification | 2 | 3.2 x 3.2 x 3.2 | 3T |
| 18 | Word identification | 2 | 3.2 x 3.2 x 3.2 | 3T |
| 19 | Object priming | 2 | 4 x 4 x 4 | 3T |
| 20 | Object retrieval | 2 | 4 x 4 x 4 | 3T |
| 21 | Object shuffle | 2 | 3.2 x 3.2 x 3.2 | 3T |
| 22 | Resting state | 2.5 | 3 x 3 x 3.5 | 3T |

**Supp. Table 3:** All *Neurosynth* term associations for CO1 regions. *Note: Numbers assigned to CO region names correspond to the ordering of CO regions as in Dworetsky* et al.*, 2021; coordinates are displayed in MNI space. Italicized gray items represent the anatomically-related terms excluded from the word clouds in* ***Figure 6.***

| CO6 | CO5 | CO15 | CO11 | CO14 | CO13 | CO12 | CO4 |
|---|---|---|---|---|---|---|---|
| **7, 8, 51** | **-3, 6, 53** | **36, 22, 3** | **-36, 20, 3** | **6, 22, 28** | **5, 20, 37** | **-1, 15, 44** | **-5, 18, 34** |
| *supplementary* | motor | *anterior insula* | *anterior insula* | *anterior cingulate* | *anterior cingulate* | task | pain |
| *supplementary motor* | *supplementary* | *insula* | *insula* | *cingulate* | *cingulate* | working memory | *cingulate* |
| *pre sma* | *supplementary motor* | task | *insular* | pain | *anterior* | working | *anterior cingulate* |
| motor | *premotor* | gain | *insula anterior* | *anterior* | *cingulate cortex* | tasks | *anterior* |
| *premotor* | tasks | *insular* | *anterior insular* | *cingulate cortex* | pain | *parietal cortex* | *cingulate cortex* |
| task | task | *anterior insular* | gain | *insula* | *acc* | *pre sma* | *anterior insula* |
| *pre supplementary* | *premotor cortex* | *anterior* | pain | *anterior insula* | *dorsal anterior* | *frontal* | *insula* |
| *motor pre* | *pre sma* | mood | *anterior* | *acc* | gain | *medial frontal* | *acc* |
| eye fields | complexity | tasks | painful | *insula anterior* | painful | performance | painful |
| movements | preparation | *insula anterior* | *inferior frontal* | *dorsal anterior* | *anterior insula* | conflict | *dorsal anterior* |
| tasks | working | working memory | *frontal* | *cortex acc* | *cortex acc* | *anterior cingulate* | noxious |
| *cortex supplementary* | execution | working | *anterior cingulate* | experiencing | task | verbal | *supplementary* |
| motor imagery | working memory | calculation | *insular cortex* | *dacc* | conflict | demands | *supplementary motor* |
| *premotor cortex* | *parietal* | *insular cortex* | *inferior frontal* | *salience network* | monitoring | mood | *cortex supplementary* |
| control | load | short term | load | competition | inhibit | *parietal* | *cortex acc* |
| *primary motor* | *frontal* | *anterior cingulate* | *acc* | painful | executive | *frontal cortex* | mood |
| eye | *motor sma* | demands | *cortex anterior* | gain | noxious | *acc* | *insula anterior* |
| execution | *motor pre* | pain | mood | risk taking | oddball | memory | *midbrain* |
| *parietal* | production | maintenance | task | sustained attention | effortful | gain | |
| *motor cortex* | *motor cortex* | *intraparietal sulcus* | *ifg* | regulatory | error | memory task | |
| *motor sma* | reading | painful | *medial frontal* | *anterior insular* | mood | phonological | |

| | | | | | | |
|---|---|---|---|---|---|---|
| *fronto parietal* | demands | difficulty | interference | empathic | *dorsolateral prefrontal* | memory wm |
| *frontal* | eye fields | *intraparietal* | *frontal gyrus* | conflict | inhibition | verbal fluency |
| *anterior insula* | language | *pre sma* | *insula inferior* | executive | *insula* | monitoring |
| | words | modality | *gyrus ifg* | noxious | frontal eye | word |
| | finger | load | demands | *somatosensory cortices* | stroop | *cingulate* |
| | *primary motor* | orthographic | *broca* | somatosensory | attentional | *cortex acc* |
| | motor imagery | risk taking | *cingulate* | | *dorsolateral* | english |
| | phonological | cognitive control | tasks | | working | *cingulate cortex* |
| | sequence | *dorsolateral* | nociceptive | | working memory | ptsd |
| | abstract | phonological | *prefrontal* | | *supplementary* | calculation |
| | *dorsal premotor* | *frontal* | | | *prefrontal* | task difficulty |
| | words | *secondary somatosensory* | | | | *anterior* |
| | movement | *parietal* | | | | motor |
| | generation | *prefrontal* | | | | |
| | interference | | | | | |
| | lexical | | | | | |
| | *cerebellum* | | | | | |
| | orthographic | | | | | |
| | speech production | | | | | |
| | finger movements | | | | | |
| | verb | | | | | |
| | action | | | | | |
| | *pre supplementary* | | | | | |
| | frontal eye | | | | | |
| | preparatory | | | | | |
| | movements | | | | | |

**Supp. Table 4:** All *Neurosynth* term associations for CO2 regions. *Note: Numbers assigned to CO region names correspond to the ordering of CO regions as in Dworetsky* et al.*, 2021; coordinates are displayed in MNI space. Italicized gray items represent the anatomically-related terms excluded from the word clouds in* ***Figure 6.***

| CO8 | CO9 | CO7 | CO3 | CO2 | CO1 |
|---|---|---|---|---|---|
| **37, 4, -4** | **49, 8, -1** | **36, 10, 1** | **-34, 3, 4** | **-45, 3, 9** | **-51, 8, -2** |
| *insula* | *insula* | *insula* | *insula* | motor | motor |
| *posterior insula* | motor | *anterior insula* | pain | *insula* | *premotor* |
| noxious | painful | *insular* | painful | muscle | imagery |
| painful | *supplementary motor* | *putamen* | *insular* | pain | phonological |
| pain | painful | pain | nociceptive | *primary motor* | *supplementary motor* |
| taste | *supplementary* | noxious | *putamen* | movement | motor imagery |
| *insular* | *operculum* | painful | *posterior insula* | *motor sma* | *somatosensory cortices* |
| *anterior insula* | externally | sensation | somatosensory | *supplementary motor* | *supplementary* |
| *insular cortex* | *insula anterior* | *caudate nucleus* | *anterior insula* | *supplementary* | *premotor cortex* |
| *putamen* | force | *posterior insula* | affective | *motor network* | finger |
| sensations | *sensorimotor cortex* | *cingulate* | *insular cortex* | *motor cortex* | *insula* |
| *somatosensory cortices* | *motor sma* | *anterior* | *basal ganglia* | finger tapping | visual word |
| autonomic | *cerebellum* | *insular cortex* | *ganglia* | *m1* | *primary motor* |
| *orbitofrontal* | *posterior insula* | *caudate* | *basal* | *premotor* | movement |
| *secondary somatosensory* | movement | *nucleus* | *somatosensory cortex* | finger | coordination |
| | *anterior insular* | *primary* | motor | painful | motor task |
| | *primary motor* | *primary secondary* | *primary* | finger movements | somatosensory |
| | *anterior insula* | arousal | interoceptive | *putamen* | *motor cortex* |
| | motor performance | risk taking | *secondary somatosensory* | autonomic | speaking |
| | *insular* | autonomic | *insula anterior* | noxious | *anterior superior* |
| | *thalamus* | *anterior cingulate* | force | movements | *hemisphere* |
| | finger | *secondary somatosensory* | *primary motor* | electrical | production |
| | *insula cortex* | *insula anterior* | *contralateral* | *ventral premotor* | movements |
| | *anterior* | *striatum* | *limbic* | *premotor cortex* | word form |
| | *primary* | *orbitofrontal* | intensity | *subcortical* | speech |
| | noxious | | *anterior insular* | tapping | painful |
| | *premotor* | | *primary somatosensory* | *cortex m1* | *ipsilateral* |
| | somatosensory | | *thalamus* | *thalamus* | *primary* |
| | | | *striatal* | aphasia | *contralateral* |
| | | | *amygdala* | *secondary somatosensory* | *cerebellum* |

| | stimulation | arm | speech production |
|---|---|---|---|
| | *somatosensory cortices* | imitation | language |
| | *supplementary motor* | execution | |
| | | *cerebellum* | |
| | | force | |
| | | motor imagery | |
| | | *somatosensory cortices* | |
| | | hand | |
| | | somatosensory | |
| | | *primary* | |
| | | *ipsilateral* | |
| | | *sensorimotor cortex* | |

### SUPPLEMENTAL FIGURES

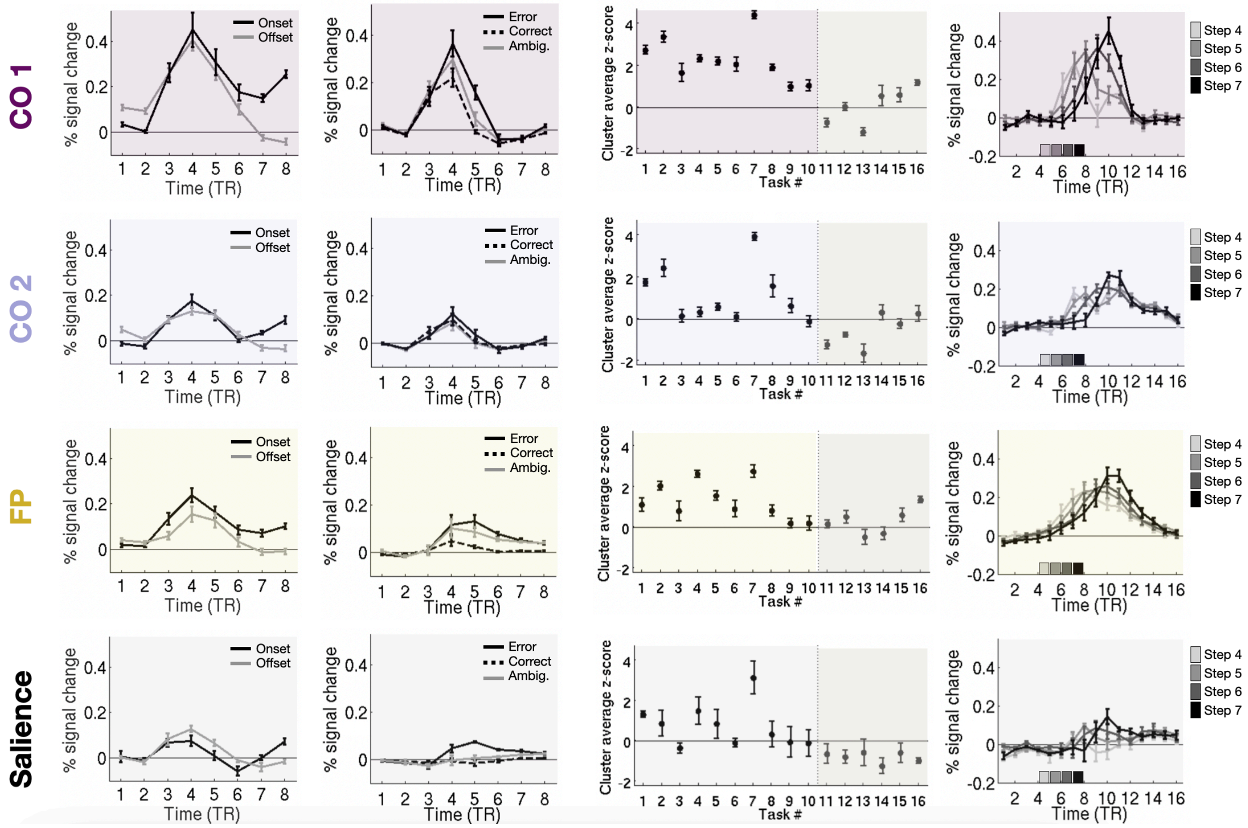


**Supp. Fig. 1**: All signals by network cluster. Task control signals (columns) are shown separated by network (rows). The plot includes signals from task initiation (onset and offset cues), task adjustment (responses to ambiguous stimuli, error, and during decision making contexts), and sustained task signals. The CO1 and CO2 clusters differ in these signals relative to each other and other large-scale networks including the frontoparietal (FP) and salience network. The CO1 sub-system shows the strongest task control characteristics of all three types.

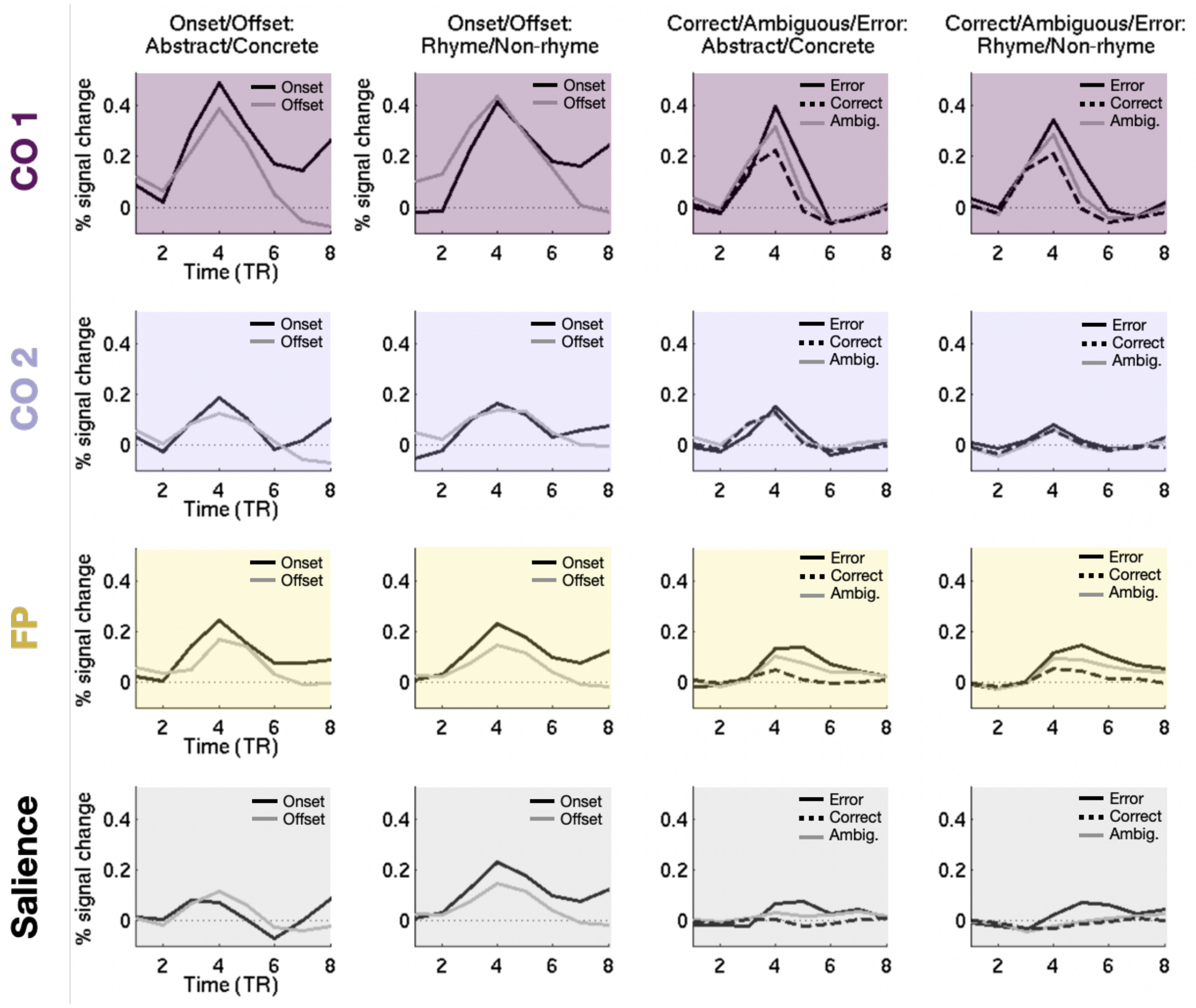


**Supp. Fig. 2**: Control signals per task. Timecourses related to task set initiation (onset/offset) and performance adjustment (correct/ambiguous/error), are shown separated by network (rows) and tasks (columns) . The two CO sub-systems are shown separately (CO1 in dark purple, CO2 in light purple) as are responses from the frontoparietal (FP) and salience networks. As can be seen, control-related responses were most robust in CO1, with strong onset and offset responses, and differences between error and ambiguous trials vs. correct; these characteristics replicated across tasks. The CO1, frontoparietal, and salience network responses for these signals were more modest, also replicating across tasks

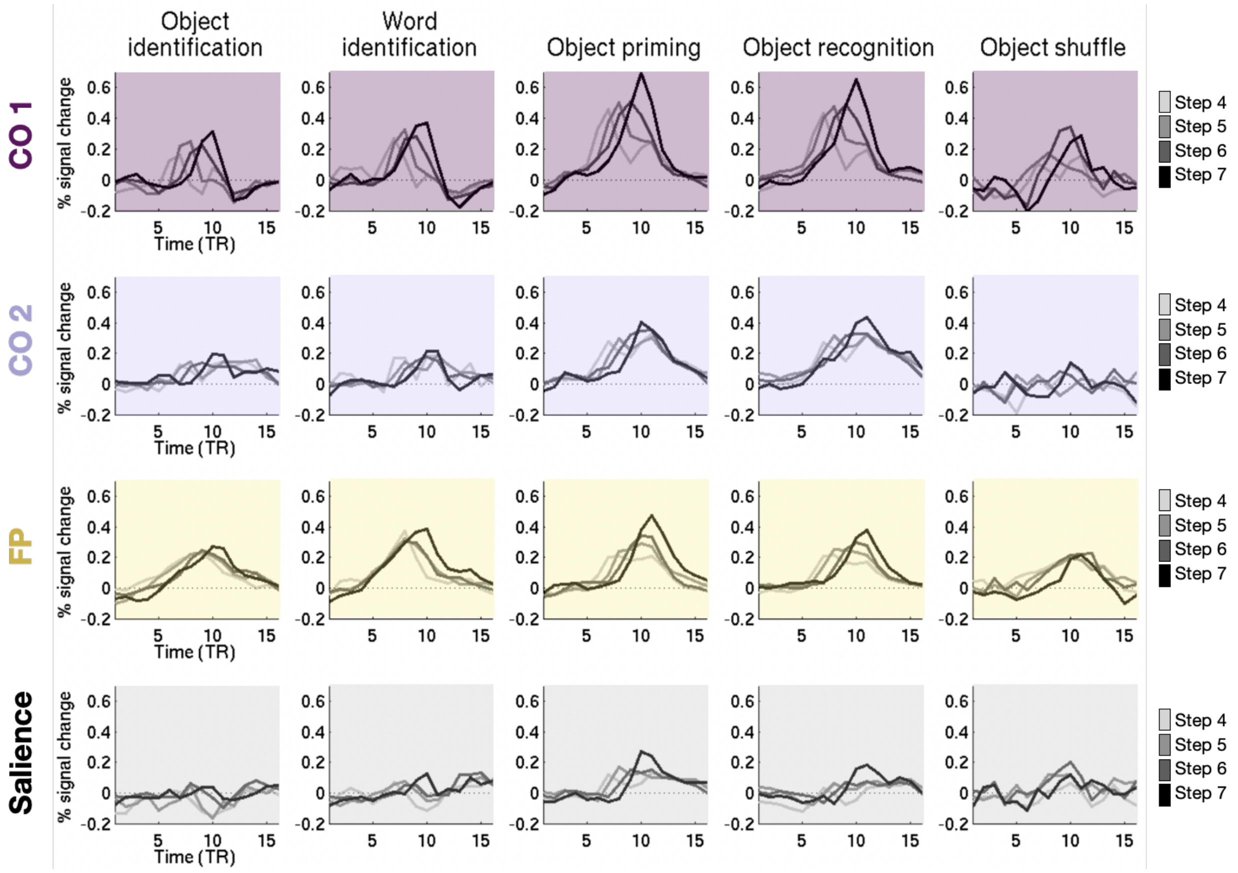


**Supp. Fig 3:** Slow reveal task signals by task. Timecourses for the slow-reveal decision making task are shown separated by network (rows) and tasks (columns). In each task, the CO1 and CO2 clusters looked clearly distinct from one another. The CO1 cluster had transient responses associated with the moment of decision across all tasks, although with variability in their magnitude. The frontoparietal cluster had early onset and graded responses peaking around the moment of decision in most tasks. The CO2 and salience clusters had dampened and more variable responses across tasks.

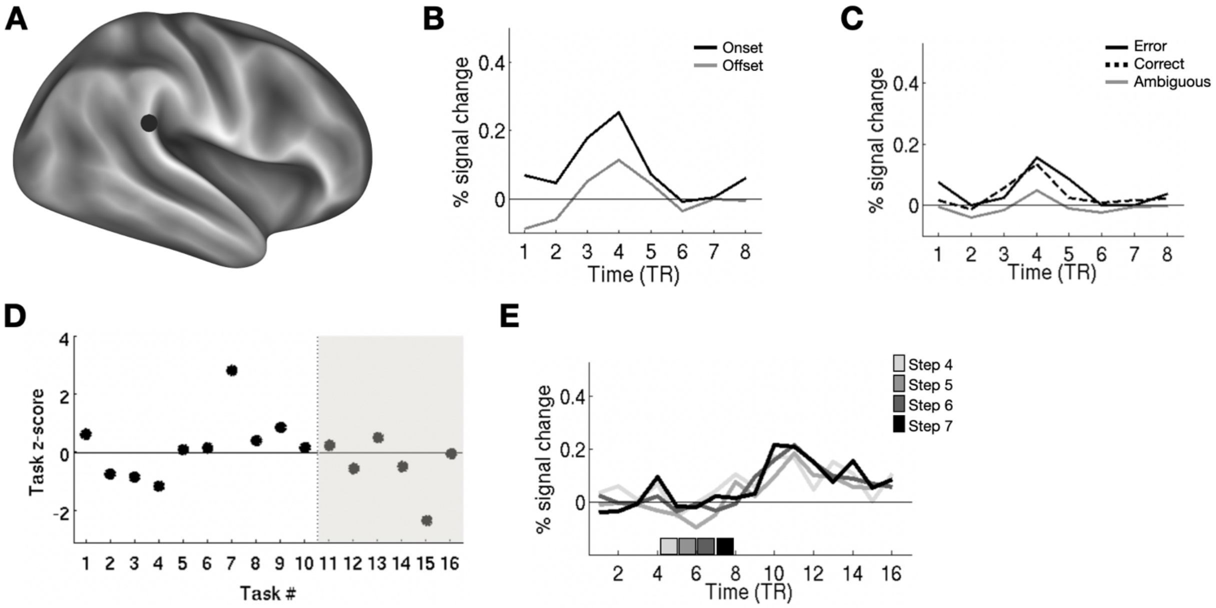


**Supp. Fig. 4:** Task signals for CO region #10, which did not distinctly cluster with the CO1 or CO2 regions. **(A)** Location on the brain, in black; **(B)** Onset vs. offset timecourses; **(C)** Correct vs. error vs. ambiguous trial timecourses; **(D)** Sustained task z-scores; **(E)** Slow reveal task timecourses.This region had relatively weak onset and offset cues, and weak responses to correct trials and errors (as in CO2). Ambiguous stimulus responses were below the level seen for correct trials. Sustained signals were very low or negative for all but one task (below the levels seen for either CO1 or CO2), and decision-making responses were weak, noisy, and ungraded regardless of the moment of decision. Thus, this outlier region did not demonstrate evidence of task control activity.

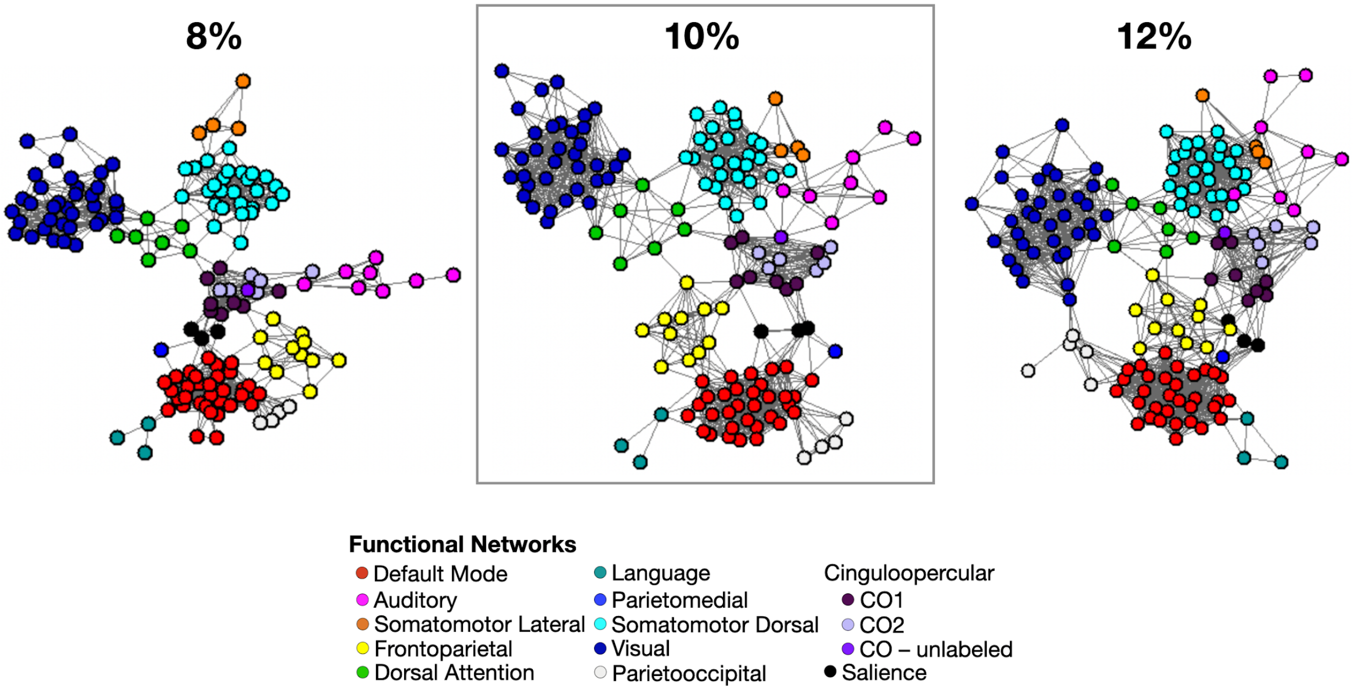


**Supp. Fig. 5:** Spring-embedded plots depicting the relationship among high consensus regions from across the whole brain, similar to that shown in **Fig. 5**. Here, we show spring embedding plots at 8% and 12% edge densities as well as the 10% edge density shown in the main text. In all cases, the CO network regions (purples) cluster together relative to regions in other networks. The CO1 sub-system is positioned closer to other putative control networks (e.g., frontoparietal, salience) relative to the CO2 sub-system.

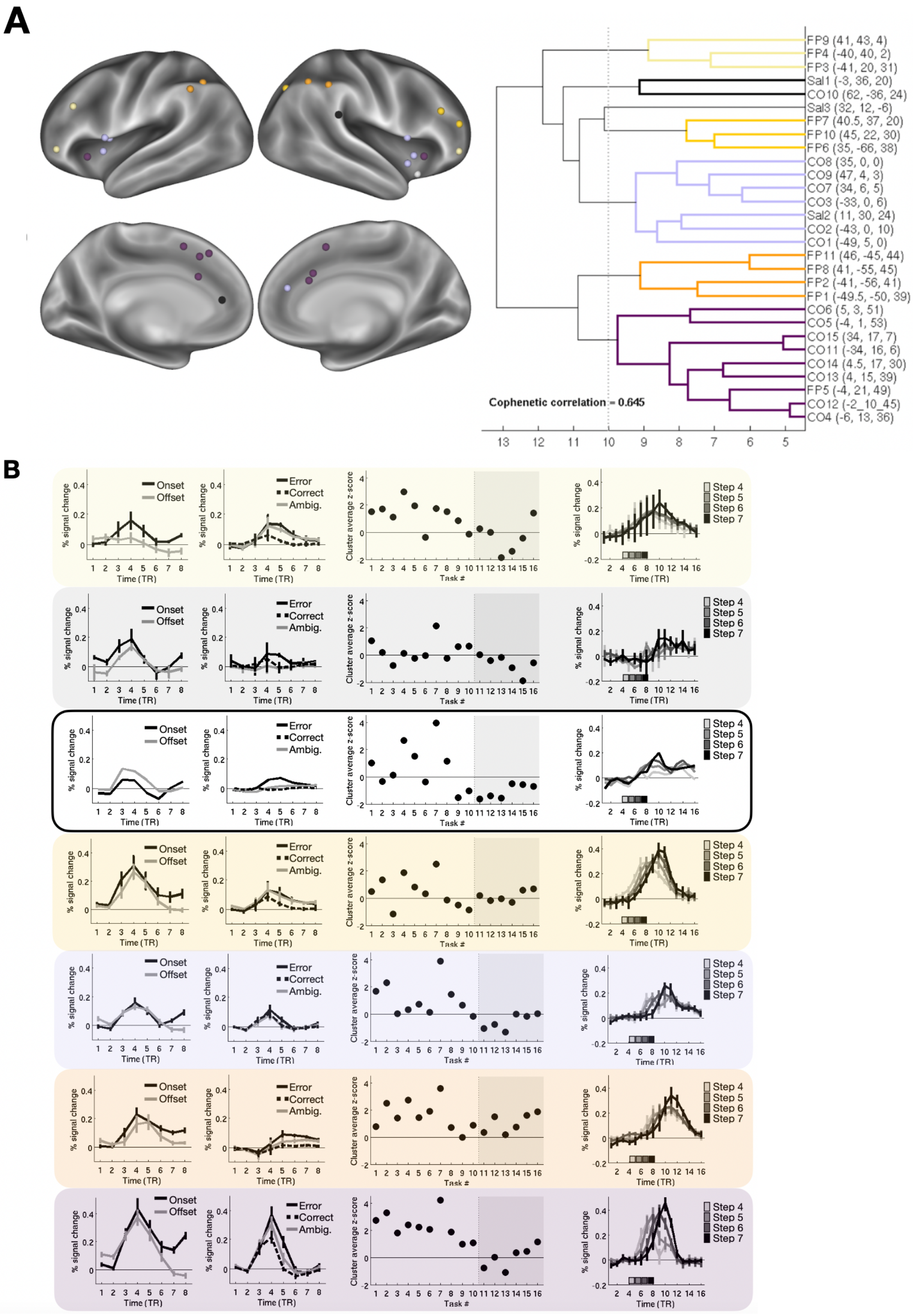
A
FP9 (41, 43, 4)
FP4 (-40, 40, 2)
FP3 (-41, 20, 31)
Sal1 (-3, 36, 20)
CO10 (62, -36, 24)
Sal3 (32, 12, -6)
FP7 (40.5, 37, 20)
FP10 (45, 22, 30)
FP6 (35, -66, 38)
CO8 (35, 0, 0)
CO9 (47, 4, 3)
CO7 (34, 6, 5)
CO3 (-33, 0, 6)
Sal2 (11, 30, 24)
CO2 (-43, 0, 10)
CO1 (-49, 5, 0)
FP11 (46, -45, 44)
FP8 (41, -55, 45)
FP2 (-41, -56, 41)
FP1 (-49.5, -50, 39)
CO6 (5, 3, 51)
CO5 (-4, 1, 53)
CO15 (34, 17, 7)
CO11 (-34, 16, 6)
CO14 (4.5, 17, 30)
CO13 (4, 15, 39)
FP5 (-4, 21, 49)
CO12 (-2_10_45)
CO4 (-6, 13, 36)
Cophenetic correlation = 0.645
13 12 11 10 9 8 7 6 5
B
% signal change
Time (TR)
Onset
Offset
Error
Correct
Ambig.
Cluster average z-score
Task #
Step 4
Step 5
Step 6
Step 7

**Supp. Fig. 6:** Hierarchical analysis is repeated with regions from all 3 networks. **(A)** Hierarchical clustering of frontoparietal (FP), salience (Sal), and cingulo-opercular (CO) regions based on task control signals, as in **Fig. 2**. Regions are shown on the left, colored by their clusters, shown on the right. **(B)** Control signals for each cluster, shown with the same cluster colors. As can be seen, similar CO1 and CO2 clusters are identified even within this broader set of regions. When the frontoparietal and salience regions are included, one frontoparietal region (FP5) is included in the CO1 cluster and one Salience region (Sal2) is included in the CO2 cluster. The remaining FP regions cluster into three distinct sub-systems (light yellow, marigold, and orange). The remaining Salience regions are included in the black cluster or singly. The remaining FP regions separate into 3 major sub-systems (FP1 in light yellow, FP2 in marigold, and FP3 in orange). The FP3 cluster (with several regions along the inferior parietal lobule) shows the strongest control-related response characteristics, most closely clustered with CO1; however, in this group, cueing responses and especially responses to error, correct, and ambiguous are diminished relative to CO1. The FP2 cluster (with regions along lateral prefrontal cortex) has an intermediate profile, with strong cueing and decision-making responses, modest/weak sustained responses, and weak (but differentiable) responses to errors and ambiguous signals relative to correct trials. The remaining salience region couples with the single supramarginal gyrus CO region and shows a similar profile shown in **Supp. Fig. 3**. Thus, there are clear differences in control task responses across these three networks, although these differences appear more subtle than what is seen in the functional connectivity data.
